# HIF2α inhibits glutaminase clustering in mitochondria to sustain growth of clear cell Renal Cell Carcinoma

**DOI:** 10.1101/2024.05.04.592520

**Authors:** Wencao Zhao, Boyoung Kim, Nathan J Coffey, Schuyler Bowers, Yanqing Jiang, Caitlyn E. Bowman, Michael Noji, Cholsoon Jang, M. Celeste Simon, Zoltan Arany, Boa Kim

## Abstract

Clear cell renal cell carcinomas (ccRCC) are largely driven by HIF2α and are avid consumers of glutamine. However, inhibitors of glutaminase1 (GLS1), the first step in glutaminolysis, have not shown benefit in phase III trials, and HIF2α inhibition, recently FDA-approved for treatment of ccRCC, shows great but incomplete benefits, underscoring the need to better understand the roles of glutamine and HIF2α in ccRCC. Here, we report that glutamine deprivation rapidly redistributes GLS1 into isolated clusters within mitochondria across diverse cell types, but not in ccRCC. GLS1 clustering is rapid (1-3 hours) and reversible, is specifically driven by reduced intracellular glutamate, and is mediated by mitochondrial fission. Clustered GLS1 markedly enhances glutaminase activity and promotes cell death under glutamine-deprived conditions. HIF2α prevents GLS1 clustering, independently of its transcriptional activity, thereby protecting ccRCC cells from cell death induced by glutamine deprivation. Reversing this protection, by genetic expression of GLS1 mutants that constitutively cluster, enhances ccRCC cell death in culture and suppresses ccRCC growth *in vivo*. These findings provide multiple insights into cellular glutamine handling, including a novel metabolic pathway by which HIF2α promotes ccRCC, and reveals a potential therapeutic avenue to synergize with HIF2α inhibition in the treatment of ccRCC.

## Introduction

Glutamine is the most abundant amino acid in human blood and is crucial for cellular growth and survival^1–3^. Glutamine is an important anaplerotic source of carbons for the tricarboxylic acid (TCA) cycle and a nitrogen source for various processes, thus contributing significantly to both ATP production and biomass synthesis in numerous cell types^2,4–6^. Glutamine is also essential for the synthesis of non-essential amino acids (NEAAs) such as asparagine, provides the backbone for glutathione in most cells, and is an important gluconeogenic precursor and regulator of urinary pH in the kidney^6^. Depletion of glutamine induces cell death across diverse cell types, variably attributable to energy depletion, inhibition of the mTOR pathway, or the initiation of endoplasmic reticulum (ER) stress^4^. The tumor microenvironment is often nutrient-poor due to reduced vascularization and/or the hyperactive metabolism of cancer cells. Glutamine has been demonstrated by many studies^7–11^, though not all^12,13^, to be one of the most depleted metabolites in tumors compared to corresponding normal tissues. Consistent with this fact, tumors are generally avid consumers of glutamine, thus creating a potential liability.

Glutaminase (GLS1) enzyme mediates the initial step in glutamine catabolism, the conversion of glutamine to glutamate, releasing a single ammonium ion^14^. There are two isozymes of GLS, GLS1 and 2, encoded by separate genes^15,16^. GLS2 expression is largely restricted to periportal hepatocytes, where the enzymes couple ammonia liberation to the production of urea^17^. GLS1 is expressed in most tissues and cancer, and encodes two alternatively spliced isoforms, kidney glutaminase A (KGA), expressed largely in kidney, and the widely expressed and more active glutaminase isoform C (GAC)^16^. All GLS1 enzymes are localized in the mitochondrial matrix^16^. GLS1 is generally thought to be regulated by tetramerization from an inactive dimer, a process requiring inorganic phosphate^18,19^. More recent work has shown that GLS1 can further oligomerize into large filamentous structures with additional enhancement of enzymatic activity^20–23^. The biological relevance of these findings is poorly understood.

Clear cell renal carcinoma (ccRCC) accounts for ∼80% of renal malignancies^24^ with a 5-year survival rate of ∼50%, but reduced to ∼10% when metastatic^25^. ccRCC is usually driven by the genetic or epigenetic loss of pVHL function^24^. In familial cases of ccRCC, there is heterozygous inheritance of *VHL* mutations, with loss of heterozygosity in the tumors^26^. Like many tumors, ccRCC tumors are avid consumers of glutamine, and glutaminase has thus long been entertained as a possible therapy for ccRCC. However, despite some efficiency with glutaminase inhibition in preclinical models, glutaminase inhibition was not effective in a recent phase 3 placebo-controlled, double blind, randomized clinical trial (PCDB-RCT)^27^. pVHL is a component of an E3 ubiquitin ligase complex required for the degradation of HIFs in the presence of oxygen. Loss of *VHL* in ccRCC is associated with stabilization of HIF, and HIF2a is likely the most important driver of ccRCC. Inhibition of HIF2α with belzufitan, a PT2385 analog, was approved for treatment of familial ccRCC in 2021, after a phase 2, open-label study showed activity in patients with ccRCC, representing a first-in-class drug approval^28^. The approval was expanded to all ccRCC in 2023 after a phase III PCDB-RCT showed marked improvements over everolimus^29^. However, although impressive, responses in both studies were largely partial, and seen only in ∼50% of patients^30^. There is thus an urgent need to better understand both the handling of glutamine and the mechanisms of HIF2α action in ccRCC.

In this study, we investigated novel molecular and cellular mechanisms of GLS1 regulation and their impact in ccRCC biology. While studying the effects of glutamine deprivation on endothelial cells, we noted dramatic clustering of GLS1 to discrete punctae throughout the cells. We used a range of pharmacological and genetic approaches to examine this process in depth, including its kinetics, its mechanism, and its impact on cellular GLS1 activity. Moreover, we identified HIF2α as a key regulator of GLS1 clustering, and demonstrate that constitutive GLS1 clustering suppresses ccRCC tumor grown *in vivo*.

## Results

### Glutamine deprivation uniquely triggers the clustering of glutaminase within mitochondria

While studying the effects of glutamine deprivation (noQ) on human umbilical vein endothelial cells (HUVECs)^31^, we incidentally noted a striking redistribution of GLS1 to discrete punctae throughout the cell, as seen by immunocytochemistry and wide-field fluorescence microscopy imaging, 24-hour after noQ (Fig. 1A). Co-staining of GLS1 with markers for various intracellular organelles, including Lamp1 for lysosomes, Calnexin for endoplasmic reticulum, BODIPY dye for lipid droplets, and Golgin for the Golgi apparatus, during noQ revealed no colocalization of GLS1 punctae with these organelles (SFig 1). In contrast, confocal and Airyscan imaging and co-staining with COXIV (Fig. 1B) or MitoTracker Red (Fig. 1C), both markers of mitochondria, demonstrated the noQ-induced GLS1 punctae to represent clustering of GLS1 within mitochondria themselves. Biochemical cellular fractionation assays, coupled with western blot analysis, confirmed the persistent presence of GLS1 within mitochondria following noQ treatment (Fig. 1D). The clustering of GLS1 represented redistribution of existing GLS1 pools, rather than de novo synthesized GLS1, because treating cells with the protein synthesis inhibitor cycloheximide prior to removing glutamine did not prevent clustering (SFig 2). The phenomenon was also not restricted to HUVECs and was observed in other cell lines tested, including HeLa, 293T, HCT116, and HepG2 (SFig 3). Finally, and importantly, noQ-induced clustering of GLS1 was unique, as it did not occur with other mitochondrial proteins, including GLUD, COXIV, CS, PDH, HADHA, and COX4l1 (SFig 4).

**Figure 1.**
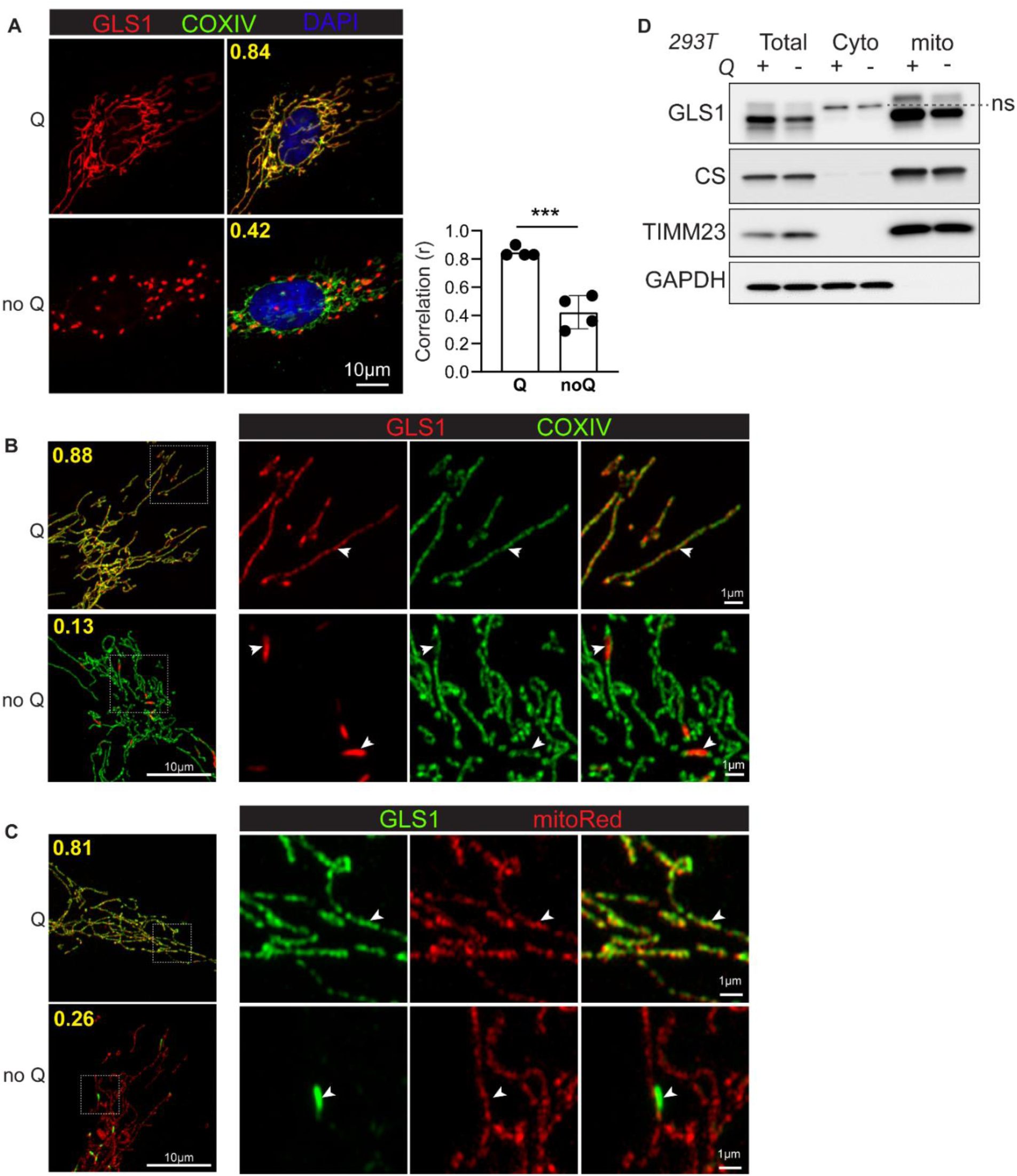
Glutamine deprivation induces GLS1 clustering within mitochondria. **A**. Immunocytochemistry (ICC) of GLS1 (red) in HUVECs after 24-hour culture in glutamine-supplemented (Q) vs. -deprived (noQ) media. GLS was co-stained with COXIV (green) and DAPI (blue). Images were taken with a wide-field fluorescence microscope using a 100x objective lens. The correlation coefficient (r) of GLS1 and COXIV staining was calculated using CellProfiler. *** p < 0.001 by t-test. **B**. ICC of GLS1 (red) and COXIV (green) after a 24-hour culture in Q vs. noQ media followed by imaging using a confocal and Airyscan microscope. **C**. Co-staining of GLS1 (green) with MitoTracker Red dye after 24-hour culture in Q vs noQ media followed by imaging using a confocal and Airyscan microscope. **D**. Mitochondrial fractionation assay performed in 293T cells after 24-hour culture in Q (+) vs. noQ (-) media.

### Glutaminase clustering is rapid and occurs within the physiological range of glutamine concentrations

To determine the kinetics of GLS1 clustering, we carried out a time-course study in C2C12 cells. GLS1 clustering was observed within 1 hour of glutamine deprivation and was complete by 6 hours (Fig. 2A). GLS1 clustering was also reversible, with similar kinetics, as replenishing glutamine led to the redistribution of clustered GLS1 as early as 1 hour (Fig. 2B). To determine the concentration of glutamine below which GLS1 clustering occurs, we performed a dose-response study in HUVECs. Treating cells with 100 μM glutamine for 6 hours promoted GLS1 clustering, while 300 μM did not (Fig. 2C). Because these cells are avid consumers of glutamine^31^, we measured glutamine concentration in the media at the end of the 5-hour incubation, revealing a remaining 55 μM and 223 μM, respectively (Fig. 2D), indicating that GLS1 clustering occurs within this range. Normal plasma concentrations of glutamine are ∼500 μM, but intratumor or for example brain interstitial concentrations are much lower, the latter ∼80 μM^1^. GLS1 clustering thus occurs in a range of glutamine concentration that can be found in tumors and other nutrient-poor settings.

**Figure 2.**
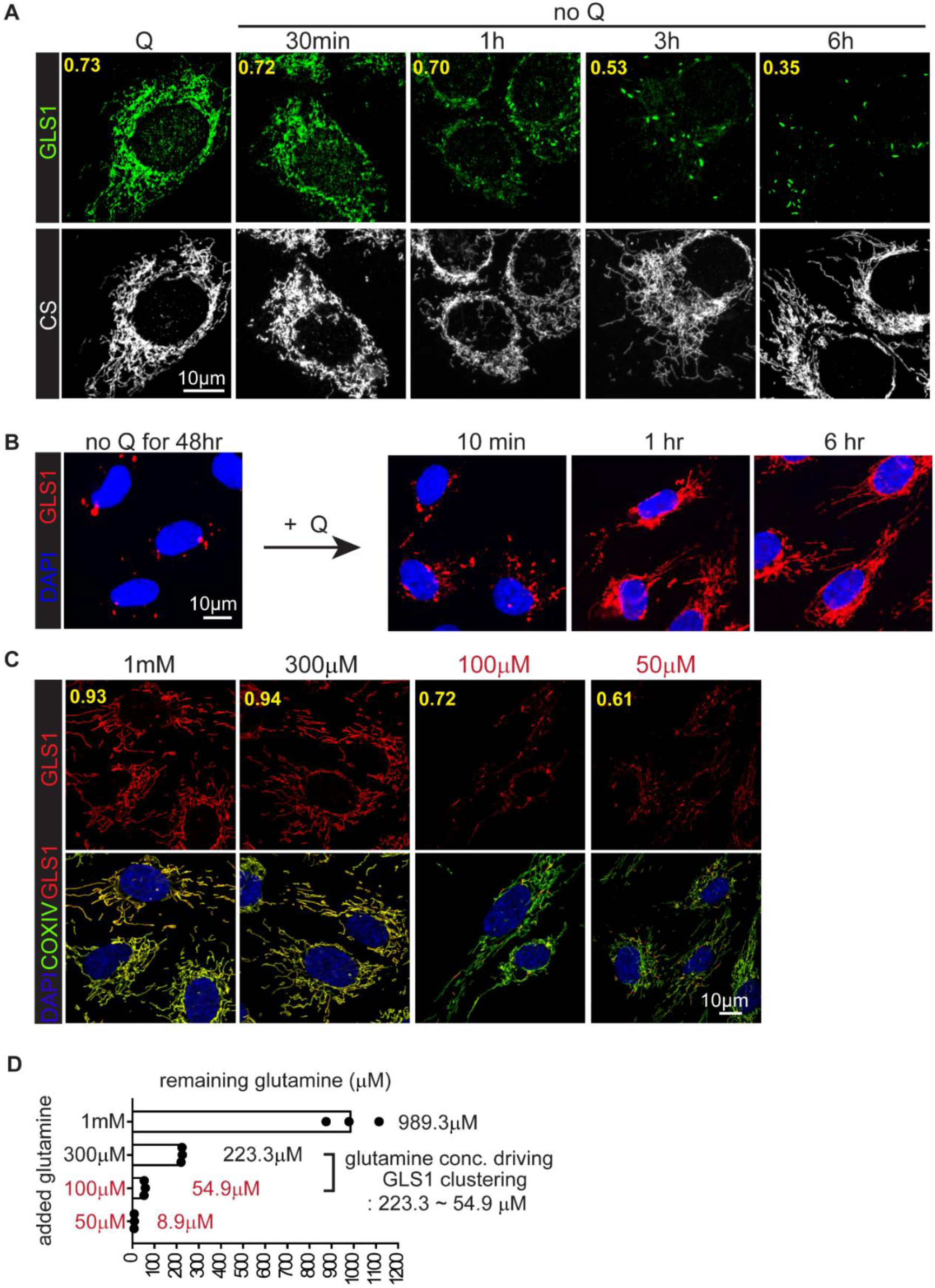
Time-course and dose-response of glutamine deprivation for GLS1 clustering. **A**. noQ time-course study in C2C12 myoblasts. Co-staining of GLS1 (green) with CS (grey) after noQ for the indicated time points. **B**. Glutamine (2mM) replenishment time course study in HUVECs after 48hrs of noQ. GLS1 (red) and DAPI. **C**. Glutamine dose-response study in HUVECs. Co-staining of GLS1 (red) with COXIV (green) after the incubation with the indicated concentrations of glutamine for 6 hours. GLS1 clustering was observed under conditions where 100μM or 50μM glutamine was incubated. **D**. Concentration of the remaining glutamine (μM) measured in the media of the experimental conditions in (C).

### Glutaminase clustering is triggered by sensing glutamate levels

We next investigated the mechanism by which GLS1 clustering is induced. We first sought to determine how the low-glutamine state is sensed. Upon cellular uptake, glutamine undergoes catabolism to glutamate by GLS1, followed by conversion to α-ketoglutarate (αKG) through the action of glutamate dehydrogenase (GLUD) and transaminases. Subsequently, αKG serves as an anaplerotic carbon source entering the TCA cycle. Conversely, αKG can contribute to the de novo synthesis of glutamate and glutamine via GLUD, transaminases, and glutamine synthase (GLUL), as illustrated in the schematic diagram (Fig. 3A). Supplementation with 2mM dimethyl-αKG, a cell-permeable form of αKG, completely inhibited GLS1 clustering (Fig. 3B-left and C). Unmethylated αKG also prevented GLS1 clustering but required a higher concentration, consistent with poor cellular uptake (SFig. 5A). We have shown previously that glutamine deprivation profoundly reduces intracellular levels of TCA intermediates, and that dimethyl-αKG largely rescues this effect^31^. Therefore, to dissect whether the rescue of GLS1 clustering by αKG relies on de novo synthesis of glutamate and glutamine versus its replenishment of TCA intermediates, we blocked αKG-to-glutamate conversion using EGCG and AOA, inhibitors of GLUD and transaminases, respectively (Fig. 3B-right and C). This treatment completely reversed the rescue of GLS1 clustering by αKG in noQ conditions, indicating that αKG-mediated rescue of GLS1 clustering is likely mediated by glutamate or glutamine, rather than replenishing TCA intermediates. Treatment with either EGCG or AOA did not fully reverse the αKG-mediated rescue of GLS1 clustering, consistent with redundancy in these pathways (SFig. 5B). αKG supplementation restored intracellular glutamate levels in noQ conditions, but did not restore glutamine levels (Fig. 3D), suggesting that glutamate, rather than glutamine, likely regulates GLS1 clustering. To confirm this conclusion, we knocked down GLUL to inhibit glutamine synthesis from glutamate, which led to further enhancement of the αKG-mediated rescue of GLS1 clustering (Fig. 3E), demonstrating that glutamate regulates GLS1 clustering independently of glutamine. Also consistent with this conclusion, supplementation with glutamate or monosodium glutamate (MSG) rescued GLS1 clustering (SFig. 5C), as did pharmacologically or genetically blocking the xCT antiporter, responsible for exporting glutamate out of the cell (SFig. 5D). Finally, considering the role of glutamine as a nitrogen source, we also tested if nitrogen depletion contributes to GLS1 clustering, but supplementing cells in noQ with ammonia showed no effect on GLS1 clustering (SFig. 5E). We conclude that GLS1 clustering is triggered specifically by sensing glutamate.

**Figure 3.**
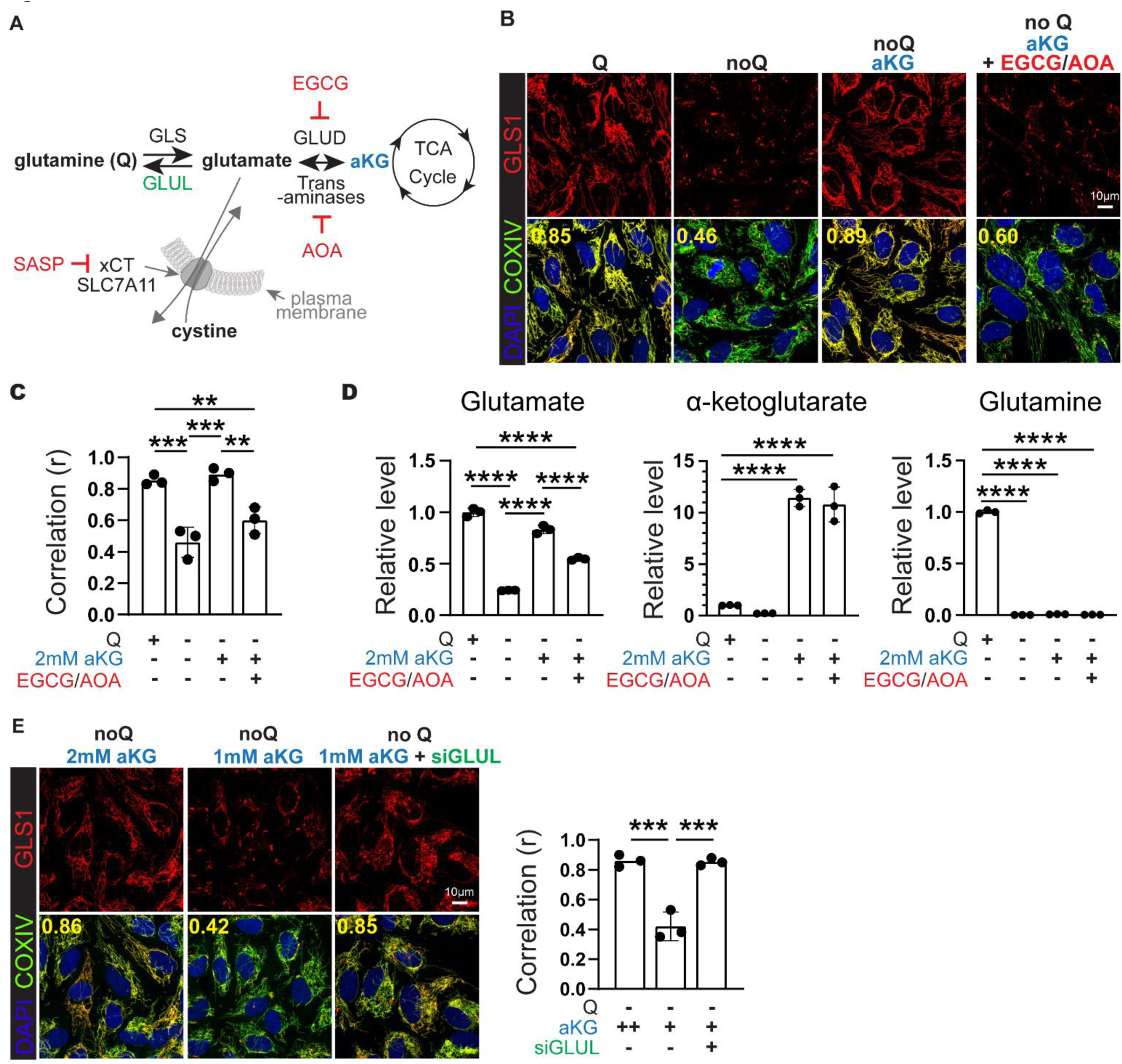
The level of glutamate, not glutamine, determines GLS1 clustering. **A**. Schematic of the rescue study, where targeted pathways and proteins are highlighted. **B**. Left: Rescue of GLS1 clustering by dimethyl-αKG (2 mM) supplementation in noQ for 6 hours. Right: Reversal of the dimethyl-αKG-mediated rescue of GLS clustering by 6-hour treatment of EGCG (100μM) and AOA (500μM) treatment. **C**. Correlation coefficient (r) for the conditions in B. *** p < 0.001, ** p < 0.01 by 1-way ANOVA. **D**. Quantifications of cellular glutamate, αKG, and glutamine in HUVECs with conditions in B. **** p < 0.0001 by 1-way ANOVA. **E**. siRNA-mediated knockdown of GLUL promotes the rescue by dimethyl-αKG (1 mM) supplementation in noQ. *** p < 0.001 by 1-way ANOVA.

### Mitochondrial fusion/fission is required for glutaminase clustering

We next sought to determine by what process the redistribution of GLS1 within the large mitochondrial network occurs. Mitochondria are highly dynamic organelles that continually undergo fusion and fission processes. The equilibrium between fusion and fission, along with significant rearrangements in the mitochondrial network, is known to vary under different metabolic states. For example, prolonged nutrient deprivation, including glutamine deprivation, promotes mitochondrial elongation^32,33^. Strikingly, however, we observed that in the short term, glutamine deprivation caused a rapid and transient fragmentation of mitochondria: mitochondrial fission was observed within an hour of glutamine deprivation (Fig. 4A), and the tubular-elongated mitochondrial network was restored within the subsequent hours, despite persistent deprivation of glutamine. Interestingly, the clustering of GLS1 in response to glutamine deprivation coincided with this transient mitochondrial fragmentation (Fig. 4B), suggesting that a fission/fusion cycle is required for the redistribution and micro-localization of GLS1 into punctae. To test this notion, we inhibited Drp1, a key effector of mitochondrial fission, either pharmacologically (mdivi-1) (Fig. 4D) or by genetic knockdown of Drp1 (Fig. 4E). In both cases, inhibition of mitochondrial fission largely prevented GLS1 clustering. We conclude that mitochondrial fission is a requisite process for GLS1 clustering in response to glutamine deprivation, likely enabling the redistribution and segregation of GLS1 within the mitochondrial network.

**Figure 4.**
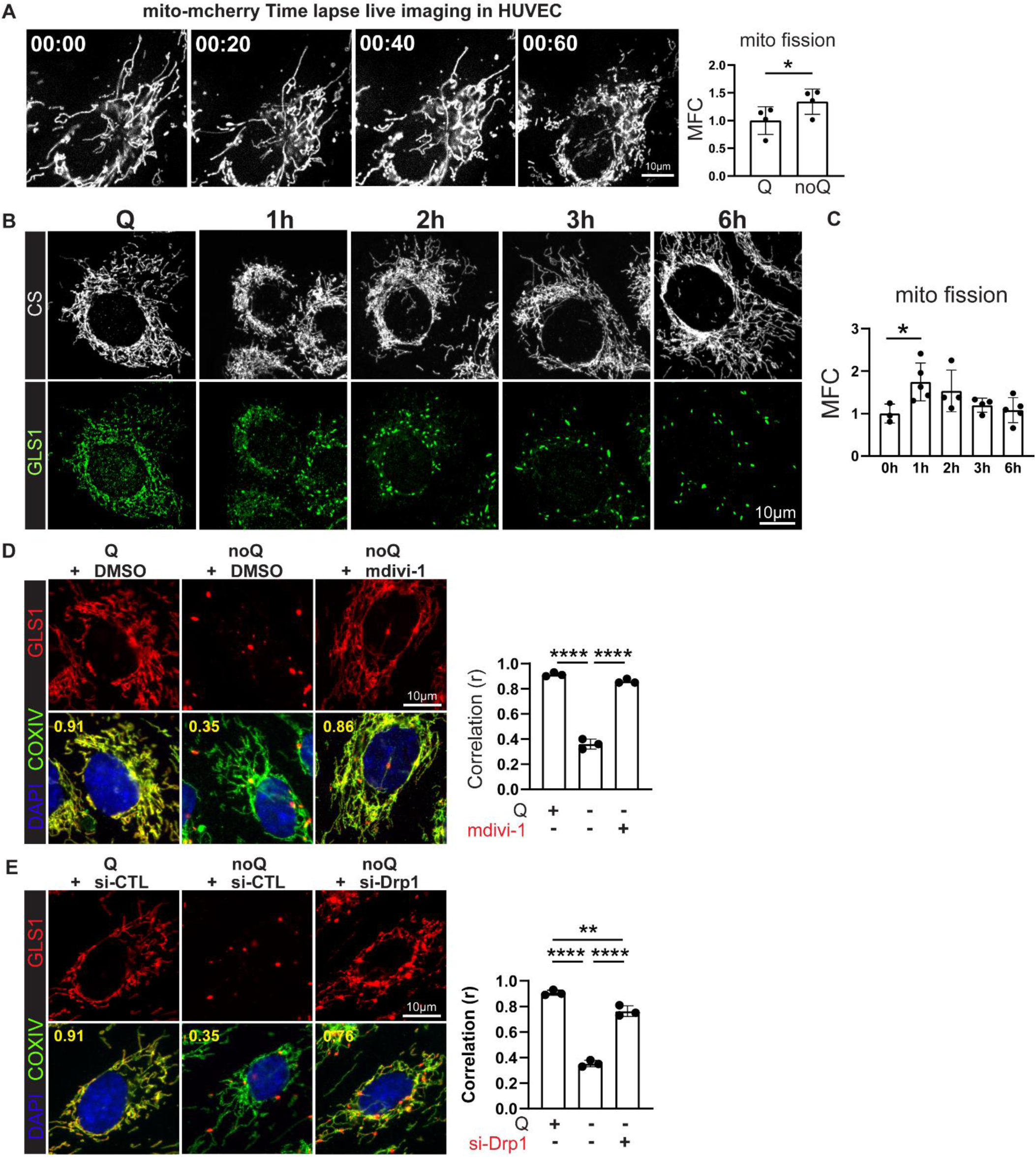
Mitochondrial fission is required for the GLS1 clustering. **A**. Timelapse live imaging of mito-mCherry overexpressing HUVECs during noQ. Quantification of mitochondrial fission is presented as mitochondrial fragmentation count (MFC). * p < 0.05 by t-test. **B-C**. ICC of CS and GLS1 during glutamine deprivation in C2C12 myoblasts. Quantification of mitochondrial fission is presented as MFC (MFC). * p < 0.05 by t-test. **D**. Rescue of GLS1 clustering by 20μM of *mdivi-1* treatment. **** p < 0.0001 by 1-way ANOVA. **E**. Rescue of GLS clustering by siRNA knockdown of Drp1. **** p < 0.0001, ** p < 0.01 by 1-way ANOVA.

### Glutaminase clustering increases its enzymatic activity

Prior biochemical studies have shown that GLS1, under some conditions such as high concentrations of inorganic phosphate, can form supratetrameric filamentous oligomers, and that this oligomerization boosted GLS1 enzyme activity^19,22^. We therefore hypothesized that GLS1 clustering in response to glutamine deprivation may similarly boost enzymatic activity, perhaps as a physiological adaptation to low substrate availability. To test this notion in intact cells, we employed a heavy isotope tracing approach. Cells were preconditioned for 24 hours in either glutamine-containing (Q) or deprived (noQ) media to induce GLS1 clustering, and subsequently additionally exposed to [U-^13^C] glutamine for 5 or 10 minutes, followed by quantification by mass spectrometry of heavy isotope-labeled glutamate, the product of the GLS1 reaction (Schematics in Fig. 5A). As shown in Fig5B-C, after 24hrs of glutamine deprivation, and thus in the context of GLS1 clustering, the amount of ^13^C-labeled glutamate (M+5) produced in 5 and 10 minutfes was more than 10 times that seen in cells maintained in complete media. Clustering in response to glutamine deprivation thus dramatically increases GLS1 enzymatic activity. The previous report indicating that GLS1 can form supratetrameric oligomers with enhanced enzymatic activity identified a critical lysine (K) at position 320, mutation of which led to constitutive oligomerization^19,22^. Consistent with this, we find that expression of this K320A mutant GLS1 (Fig. 5D) in intact cells leads to constitutive clustering of GLS1, even under glutamine-rich conditions (Fig. 5E), and concomitant markedly higher GLS1 activity than in cells expressing wild-type (WT) GLS1, as determined by [U-^13^C] glutamine tracing (Fig. 5F). In sum, intracellular GLS1 clustering induced either by glutamine deprivation or by genetic manipulation markedly increases its enzymatic activity.

**Figure 5.**
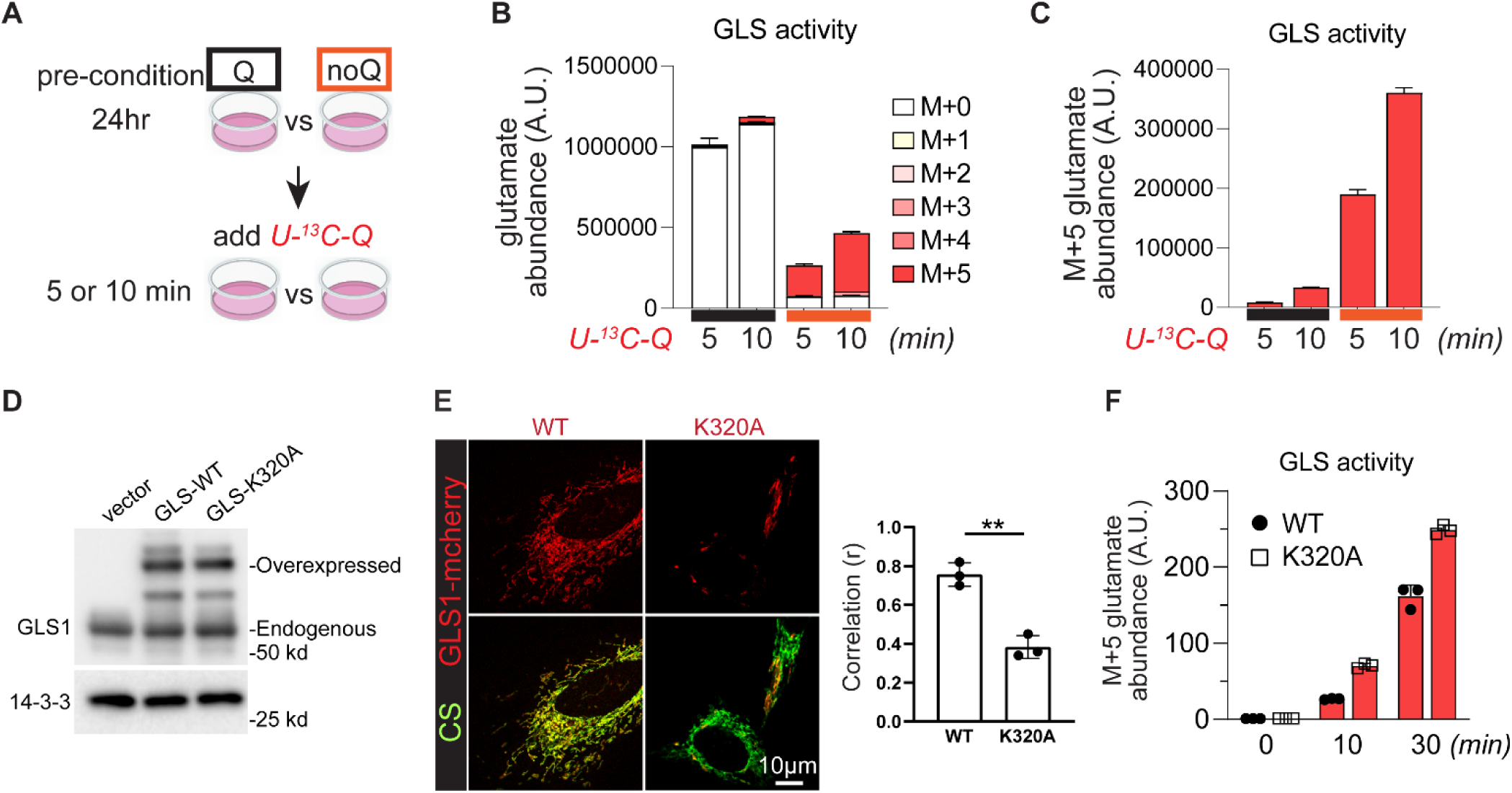
GLS1 clustering increases its enzymatic activity. **A**. Schematic of the tracing experiment using U-^13^C glutamine (Q) to measure enzymatic activity of GLS. **B**. All glutamate that are heavy isotope-labeled by the indicated number of carbons. M+n: glutamate with n carbon atoms labeled with ^13^C. **C**. Separate plot of ^13^C-glutamate that are M+5. **D**. Western blotting analysis for the validation of the equivalent expression of GLS1-WT or GLS1-K320A in Hela cells. **E**. ICC of WT-vs K320A-GLS1-mCherry overexpressing HeLa cells. ** < 0.01 by t-test. **F**. ^13^C-glutamate that are M+5 in WT-vs K320A-GLS1-mCherry overexpressing HeLa cells.

### HIF2α suppresses glutaminase clustering

In further efforts to elucidate the molecular mechanisms that drive GLS1 clustering, we tested the impact of several compounds that mimic aspects of tumor biology and identified the hypoxia-mimetic dimethyloxalylglycine (DMOG) as a potent suppressor of GLS1 clustering (Fig. 6A-D). DMOG is an αKG analogue that inhibits several αKG-dependent dioxygenases, including HIF prolyl hydroxylase, which mediate degradation of HIF transcription factors in the presence of oxygen^34^. Consistent with this, DMOG treatment stabilized HIF1α and HIF2α proteins (Fig. 6E) and induced their transcriptional activity (Fig. 6F). We hypothesized that activation of HIF1α or HIF2α, the two predominant HIF factors, may suppress GLS1 clustering. To test this notion, we investigated whether the suppression of GLS1 clustering by DMOG requires HIF1α or HIF2α. Knockdown of HIF1α (validated in Fig. 6E and SFig. 6A) did not block DMOG’s suppression of GLS1 clustering (Fig. 6B and 6D). In contrast, knockdown of HIF2α (validated in Fig. 6E and SFig. 6A) abrogated the effect of DMOG (Fig. 6C and 6D). Similarly, pharmacological inhibition of HIF2α translation (validated in Fig. 6E) also prevented DMOG’s suppression of GLS1 clustering (Fig. 6C and 6D). We conclude that HIF2α, but not HIF1α, suppresses GLS1 clustering. Interestingly, we noted that inhibitors of HIF2α transcriptional activation (HIF inhibitors VII and PT2385) did not reverse DMOG’s effect (Fig. 6G), despite efficient inhibition of HIF2α transcriptional activity (Fig. 6F), suggesting that the effects of HIF2α are not dependent on HIF2α transcriptional activity. Further supporting this conclusion, overexpression of a constitutively stabilized ΔTAD HIF2α mutant that lacks transcription activation domains rescued GLS1 clustering as efficiently as did transcriptionally active HIF2α (SFig. 6B-6D). Finally, we also observed that DMOG treatment suppressed glutamine deprivation-induced cell death (Fig. 6H), indicating a pro-survival role of HIF2α under these conditions. Together, these findings demonstrate that HIF2α, but not HIF1α, suppresses GLS1 clustering, and does so in a transcriptional activity-independent manner.

**Figure 6.**
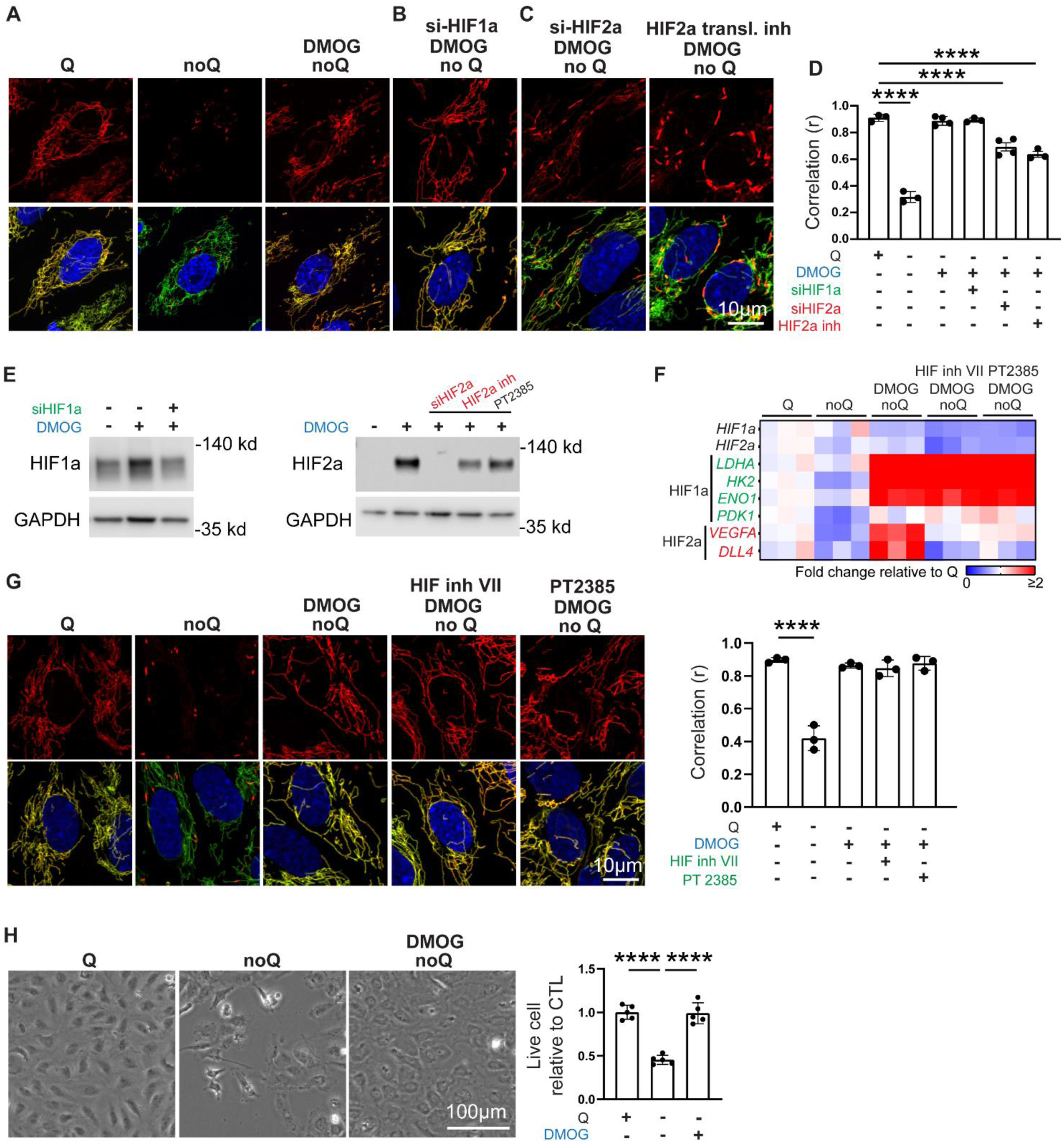
HIF2α prevents DMOG-induced redistribution of GLS1. **A**. Rescue of GLS1 clustering by DMOG treatment in HUVECs. **B**. No reversal of the DMOG-induced GLS1 redistribution by knockdown of HIF1α. **C**. Reversal of the DMOG-induced GLS1 redistribution by knockdown of HIF2α or inhibition of HIF2α translation. **D**. Correlation coefficient (r) for the conditions in **A, B** and **C**. **** p < 0.0001, ** p < 0.01 by 1-way ANOVA. **E**. Western blotting analysis showing the validation of siRNA and chemical compounds on the expression of HIF1a (left panel) or HIF2a (right panel) in HUVECs. **F**. qPCR analysis showing the validation of DMOG and HIF2a transcriptional inhibitors in HUVECs. HIF1a target genes: *LDHA*, *HK2*, *ENO1*, and *PDK1*. HIF2a target genes: *VEGFA* and *DLL4*. **G**. No reversal of the DMOG-induced GLS1 redistribution by inhibitors targeting the transcriptional activity of HIF2α. **** p < 0.0001 by 1-way ANOVA. **H**. Rescue of noQ-induced cell death by DMOG treatment in HUVECs. **** p < 0.0001 by 1-way ANOVA.

### GLS1 clustering in ccRCC is prevented by HIF2α

Activation of HIF2α is observed in nearly all cases of clear cell renal cell carcinomas (ccRCCs), usually as a result of VHL inactivation, and HIF2α is required for tumor development^35–37^. In contrast, HIF1α, while also activated by loss of VHL, is dispensable for tumor development^36,37^. We thus hypothesized that GLS1 clustering would be suppressed in HIF2α-positive ccRCC cells, and furthermore that the inhibition of GLS1 clustering by HIF2α in these cells may contribute to cell viability and tumorigenesis, as suggested above (Fig. 6H). Indeed, in contrast to all other cells we tested (Fig1, Fig2, SFig 3), both UMRC2 and 769-P cell lines, derived from human ccRCCs and which exhibit constitutive HIF2α activity^38^ (Fig. 7A-B), were resistant to GLS1 clustering under glutamine-deprived conditions (Fig. 7C-left and SFig. 7C-left). Moreover, this resistance was dependent on HIF2α as demonstrated by siRNA knockdown (Fig. 7A, 7C-right and SFig. 7A-C). Coincident with the protection from GLS1 clustering, these cells were also protected from cell death induced by glutamine deprivation, akin to the protection seen with DMOG in other cells (Fig. 6F), and suppressing HIF2α sensitized the cells to glutamine deprivation (Fig. 7D, 7E, SFig. 7D, and 7E). Conversely, ectopic expression of the constitutively clustering K320A GLS1 mutant was sufficient to induce cell death in glutamine-deprived ccRCC cell lines (Fig. 7F and SFig. 7F). These findings demonstrate that HIF2α-driven ccRCC cells actively suppress GLS1 clustering in a HIF2α-dependent fashion. Moreover, HIF2α-driven inhibition of GLS1 clustering is required to prevent cell death in low glutamine environments, uncovering an important liability of these tumors.

**Figure 7.**
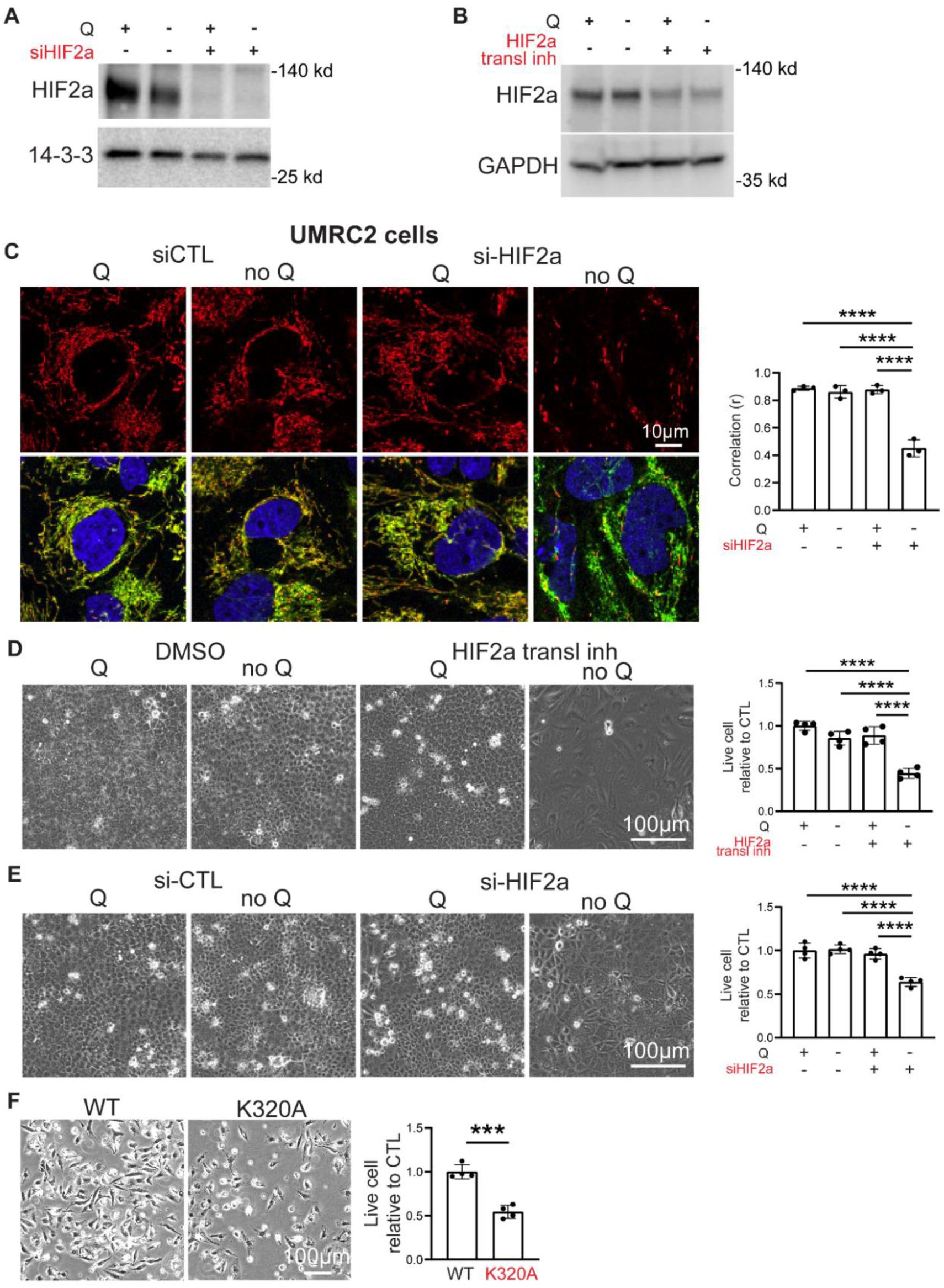
GLS1 clustering is prevented in UMRC2 cells in a HIF2α-dependent manner. **A** and **B**. Western blotting analysis for the validation of HIF2a siRNA (A) and its translational inhibitor (B) in UMRC2 cells. **C**. Resistance to noQ-induced GLS1 clustering in UMRC2 cells is reversed by si-HIF2α. **** p < 0.0001 by 1-way ANOVA. **D**. Resistance to noQ-induced cell death in UMRC2 cells is reversed by treatment with an inhibitor of HIF2α translation. **** p < 0.0001 by 1-way ANOVA. **E**. Resistance to noQ-induced cell death in UMRC2 cells is reversed by siHIF2α. **** p < 0.0001 by 1-way ANOVA. **F**. Increased cell death in UMRC2 cells overexpressing the K320A mutant GLS1 compared to those overexpressing WT GLS. *** p < 0.001 by t-test.

To investigate HIF2a-dependent inhibition of GLS1 clustering in human RCC, we performed immunohistochemistry (IHC) on primary human tissue samples, including normal kidney, ccRCC, and papillary RCC (pRCC) (SFig. 8A). We compared ccRCC and pRCC due to their distinct HIF biology: ccRCC is generally associated with VHL loss and constitutive HIF2α activation, while pRCC generally do not have HIF2α activation, instead bearing mutations in several distinct pathways^39^. Blinded immunohistochemical analyses revealed diffuse GLS1 staining seen in normal kidney samples (SFig. 8A) and clear GLS1 clustering in pRCC samples examined, likely a response to low intra-tumor glutamine concentrations. In sharp contrast, almost no GLS1 clustering was seen in ccRCC samples (only one isolated foci showed clustering), consistent with suppression of GLS1 clustering by activated HIF2α in the context of tumors lacking VHL. Together, these findings demonstrate that HIF2α-driven ccRCC cells actively inhibit GLS clustering, playing a key role in ccRCC biology, and uncover a potential liability in these lethal tumors.

### Promoting GLS1 clustering suppresses ccRCC tumor growth

To test this potential liability of GLS1 clustering in tumors *in vivo*, we used a subcutaneous xenograft model of ccRCC by subcutaneous injection of UMRC2 cells into the dorsal flanks of immunocompromised mice. Patient-derived xenografts from *VHL*-mutant ccRCC retain robust glutamine metabolism (both oxidative and reductive), supporting the relevance of the model^40^. Animals received UMRC2 cells overexpressing either WT or K320A GLS1 on paired dorsal flanks, and tumor growth was monitored noninvasively for 10 weeks, followed by sacrifice and histological analysis (Schematics in Fig. 8A). Elevated GLS activity in K320A GLS1-expressing UMRC2 cells was validated using [U-^13^C] glutamine and quantifying labeled glutamine, glutamate, and aKG (SFig. 8B-F). K320A GLS1-expressing tumors exhibited significantly reduced growth, with lower final tumor volumes and weights compared to WT controls (Fig. 8B-E). Immunohistochemistry confirmed GLS1 clustering in K320A GLS1-expressing tumors (Fig. 8F), while TUNEL staining revealed increased apoptosis in these tumors (Fig. 8G), consistent with the increased cell death observed *in vitro* (Fig. 7F and SFig7F). K320A GLS1-expressing tumors exhibited elevated glutamate levels, consistent with increased GLS1 activity *in vivo* (SFig. 8G-8H). These findings demonstrate that promoting GLS1 clustering and enhancing its activity in ccRCC tumors suppresses tumor growth and increases apoptosis.

**Figure 8.**
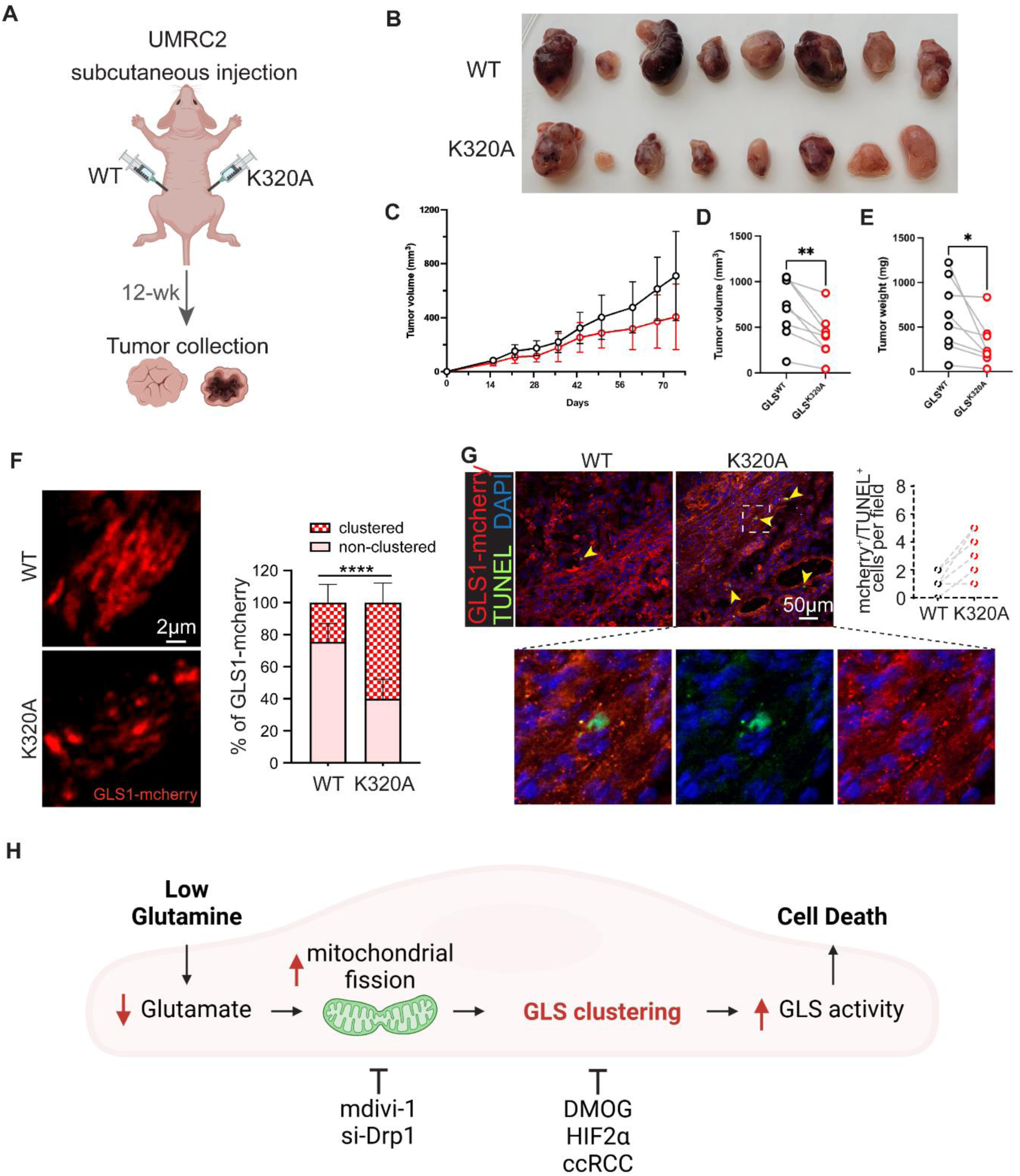
Promoting GLS1 clustering in UMRC2 suppresses tumorigenesis. **A**. Schematics of the UMRC2 injection study. Each nude mouse received subcutaneous injections of 10 million UMRC2 cells overexpressing either WT-GLS or K320A GLS1 into their left and right flanks, respectively. **B**. Reduced tumor growth in K320A-overexpressing UMRC2. Photo of all dissected tumors: WT on the upper panel and K320A on the lower panel, n=8 for each group. **C**. Average tumor volume (mm^3^) in WT-vs K320A GLS1-overexpressing UMRC2 during the 12-week monitoring period. **D**. Pair matched plot of tumor volume (mm^3^) in WT-vs K320A GLS1-overexpressing UMRC2 in each mouse. ** p < 0.01 by paired sample t-test. **E**. Plot of tumor weight (mg) in WT-vs K320A GLS-overexpressing UMRC2 in each mouse. * p < 0.05 by paired sample t-test. **F**. Confirmation of GLS1 clustering in K320A-overexpressing UMRC2 tumor. Left panel depicts the representative confocal images. Right panel shows the quantification of GLS1 clustering. **** p < 0.0001 by paired sample t-test. **G**. Increased apoptosis in K320A-overexpressing UMRC2 tumor. **H**. Schematics of the model.

## Discussion

In this study, we demonstrate that glutamine deprivation induces GLS1 clustering in various cell types, encompassing normal and cancer cells, and we elucidate the mechanisms by which GLS1 clustering occurs, as well as the enzymatic and functional consequences of GLS1 clustering. We propose a model (Fig8H) whereby low extracellular glutamine reduces intracellular glutamate levels, leading to mitochondrial fission-mediated GLS1 clustering, which increases GLS1 enzymatic activity and promotes cell death. Moreover, we show that inhibition of this process actively occurs in ccRCC and is mediated by HIF2α, elucidating a new role for HIF2α, and uncovering a liability that could be leveraged to suppress tumor growth by promoting GLS1 clustering.

As noted above, ccRCC tumors are avid consumers of glutamine, but glutaminase inhibition, long entertained as a possible therapy for ccRCC, was not effective in a recent phase 3 PCDB-RCT^27^. CB-839, the agent used in these trials, and its analog BPTES, inhibit glutaminase activity by allosteric prevention of the dimer-to-tetramer transition of GLS1^41,42^ and of GLS1 supratetrameric formation^21,22^. Here we show that, in contrast, promoting GLS1 clustering, accompanied by markedly increased GLS1 enzymatic activity, also suppresses ccRCC tumor growth in a pre-clinical xenograft model. Recent work, performed in parallel to ours, has indicated similar effects of GLS1 clustering in suppressing glioma xenograft models^23^. This raises the interesting possibility that, in certain contexts, reduced rather than increased GLS1 activity promotes ccRCC tumorigenesis. Tumors are notoriously heterogeneous, likely including areas of glutamine sufficiency but others of glutamine deprivation, where preventing GLS1 clustering may in fact be beneficial. Thus, our findings offer a possible explanation for why glutaminase inhibitors have not had therapeutic success so far.

HIF2α is likely the most important driver of ccRCC. As noted above, inhibition of HIF2α with belzufitan, a PT2385 analog, is now approved for treatment of ccRCC. However, clinical responses remain largely incomplete, and seen only in ∼50% of patients^30^, indicating the need for further understanding how HIF2α affects ccRCC biology. We now demonstrate here that HIF2α, but not HIF1α, suppresses GLS1 clustering and hyperactivation. This novel function for this oncogenic transcription factor likely critically contributes to tumor growth, because as we have shown, reversing this effect of HIF2α, i.e. constitutively forcing GLS1 clustering, reduces ccRCC tumor growth (Fig7). Understanding how mechanistically HIF2α suppresses GLS1 clustering will be of great interest. Importantly, we find that PT2385 does not prevent HIF2α from suppressing GLS1 clustering. PT2385 blocks the formation of HIF2α/ARNT heterodimers, an obligatory event for HIF2α-mediated DNA binding and transcriptional activity. Thus it appears that HIF2α blocks GLS1 clustering in a non-genomic fashion. The fact that PT2385 does not affect this function of HIF2α may in part explain the incomplete clinical efficacy of PT2385, and we predict, therefore, that there may exist an opportunity to synergize current HIF2α inhibition with additional stimulation of GLS1 clustering.

Why would too much GLS1 activity suppress tumor growth? GLS1 removes the gamma-nitrogen amide group from glutamine, converting it to glutamate and releasing an ammonium ion. The gamma-nitrogen amide group of glutamine is an indispensable donor of nitrogen for several essential cellular metabolic processes, including synthesis of asparagine, purine and pyrimidine nucleobases, and hexosamines. Glutamine is also critical for the synthesis of glutathione to defend against oxidative stress. Overactive GLS1 activity can thus be predicted, under certain circumstances, to deplete glutamine and to suppress these critical pathways. Indeed, the balance of glutamine utilization has been suggested to skew away from glutaminolysis and toward nucleotide synthesis during malignant progression^43^. We have shown before that, to survive glutamine deprivation, endothelial cells activate macropinocytosis as an alternative source of asparagine and other NEAAs normally synthesized from glutamine^31^. Additionally, Jiang *et al*. demonstrated in human embryonic brain cells that asparagine supplementation rescues cell death induced by low glutamine^23^. Thus, under some circumstances, GLS1 aggregation in response to glutamine deficiency may be a maladaptive response that skews glutamine use towards glutaminolysis, perhaps to optimizes anaplerosis, but at the expense of use of its gamma-nitrogen amide for other critical cell functions. It is also possible that the redistribution of GLS1 to specific sub-mitochondrial compartments causes metabolic channeling to specific pathways, in a way that is deleterious to cellular viability. Finally, clustered GLS1 may also have pro-apoptotic activity that is independent of its enzymatic activity. Suggestive of this possibility, Jiang *et al*. showed that αKG blocks clustering and rescues cell death (despite not providing gamma-nitrogen amide groups) in human embryonic brain cells, but αKG does not do so in cells expressing the constitutively clustered K320A mutant^23^.

How is GLS1 clustering regulated? Ferreira *et al*.^22^ and, more recently, Adamoski *et al*.^21^ using cryogenic electron microscopy (EM), demonstrated that inorganic phosphate (Pi) allosterically activates GLS1 and promotes enzyme filamentation. Adamoski *et al*^21^. further used EM to show that intra-mitochondrial GLS1 clustering seen in intact cells corresponded to GLS1 filamentation, as seen *in vitro*. To what extent Pi regulates this process under physiological conditions, however, is not clear. Glutamine deprivation does not appear to alter intracellular Pi levels^23^. Instead, we find, as do Jiang *et al*.^23^, that GLS1 polymerization is regulated by intracellular glutamate levels, within physiological ranges. Interestingly, Hans Krebs first noted in 1935 that glutaminolysis in kidney extracts (i.e. GLS1), but not liver extracts (i.e. GLS2), was exquisitely sensitive to inhibition by glutamate, and he hypothesized competitive inhibition between glutamine and glutamate^44^. Subsequent biochemical work confirmed this hypothesis, showing competition that favored glutamate binding at [Pi]=∼10mM (a typical intra-mitochondrial concentration), while favoring glutamine binding at high [Pi]^45^. Precisely how glutamate regulation occurs will require structural studies akin to those carried out by Adamoski *et al*^21^.

With respect to regulation of GLS1 clustering, we also show that glutamine deprivation triggers a transient cycle of mitochondrial fission/fusion that is required to enable clustering of GLS1 into distinct intra-mitochondrial punctae. The dynamics of mitochondrial fusion and fission are pivotal for maintaining mitochondrial quality control. Typically, mitochondrial fragmentation occurs under conditions of nutrient excess, whereas nutrient starvation induces mitochondrial elongation^46^. Consistent with this paradigm, studies have shown that glutamine deprivation leads to mitochondrial elongation within 4-6 hours^32,33^. In contrast to these observations, our study reveals a distinctive pattern: immediately after glutamine starvation, within the initial 1-2 hours, mitochondria undergo excessive fission, followed by resumption of mitochondrial networks by fusion. Prior studies may have missed this initial time window. Strikingly, we find that this rapid fission/fusion process is essential for GLS1 clustering. The fission/fusion cycle may be required to allow self-assembling GLS1 oligomers to achieve a certain size. Alternatively, the process of mitochondrial remodeling may, itself, trigger signaling that enables oligomerization, e.g. by altering intramitochondrial glutamate concentrations. Further studies of this process will be of interest.

In summary, we show here that glutamine deprivation induces GLS1 clustering in various cell types, and we reveal several mechanistic aspects of this process. Importantly, we show that HIF2α blocks GLS1 clustering in ccRCC, thereby promoting tumor growth. The work elucidates multiple aspects of glutamine handling, including a novel connection between glutamine handling and HIF2α, the predominant driver of ccRCC, thus uncovering a potential therapeutic avenue to synergize with HIF2α inhibition in the treatment of ccRCC.

## Methods

### Cell culture

293T, HeLa, HCT116, HepG2, C2C12, UMRC2 and 769-P cells were cultured in DMEM (Gibco, 11995056) supplemented with 10% FBS. HUVECs were cultured in EBM2 containing EGM supplements (Lonza, CC-3162) with 10% FBS. Glutamine deprivation studies were done using DMEM that contains no glutamine (Gibco, 31053028) supplemented with 10% dialyzed FBS (HyClone).

For siRNA and DNA transfection, cells were kept in serum-free Opti-MEM media for 6 hours of transfection duration, after which they were refreshed with their complete media. All siRNAs are treated at 10 nM concentration and are obtained from Sigma-Aldrich: human si-GLUL (SASI_Hs02_00307974), human si-SLC7A11 (SASI_Hs02_00345461), human si-DRP1 (SASI_Hs02_00340086), human si-HIF1α (SASI_Hs02_00332063), and human si-HIF2α (SASI_Hs01_00019152). For the HIF2a overexpression experiment, constitutively stabilized HIF2a (Addgene, 44027)^47^ or HIF2a ΔTAD^47^ plasmids were transfected to HUVECs. Confirmation of siRNA/DNA-mediated genetic manipulation was determined using multiple different methods including qPCR, WB or ICC.

Cell death was assessed using a commercial kit (Thermo Fisher Scientific, V13241) according to the manufacturer’s instructions. Drugs and chemicals were obtained from either Sigma or MedchemExpress (MCE) and treated at the indicated concentration in the figures: dimethyl αKG (Sigma, 349631), NH_4_Cl (Sigma, A9434), L-glutamate (Sigma, G8415), monosodium glutamate (MSG) (Sigma, G5889), cyclohexamide (Sigma 239765) (2μg/ml), EGCG (Sigma, E4143), AOA (Sigma, C13408), SASP (Sigma, S0883), mdivi-1 (20uM) (Sigma, M0199), DMOG (Sigma, D3695), HIF2α translation inhibitor (Sigma, 400087), HIF inhibitor VII (Sigma, 5043790001), and PT2385 (MCE, HY-12867).

For the isotope tracer assay using U-^13^C_5_ glutamine, UMRC2 cells were grown in 6-well plates and incubated with the growth medium (DMEM without glutamine, Gibco, 31053028) containing U-^13^C_5_ glutamine (Cambridge Isotope Laboratories, CLM-1822-H-0.1, 2 mM), 10% dialyzed FBS (Cytiva, SH30079.02) and 1% Penicillin/Streptomycin. After incubation, culture media were harvested, and cells were washed with 37°C saline and lysed immediately for LC/MS.

### Immunocytochemistry (ICC)

Cells were plated onto glass coverslips and subjected to siRNA transfection, drug treatment, and/or glutamine starvation as indicated in the figures. Cells were then fixed with 3.7% paraformaldehyde, washed, and permeabilized with 0.3% Triton X-100. After blocking with 2% BSA, samples were incubated with primary antibodies overnight. Primary antibodies used for the ICC are GLS1 (abcam, 156876), COXIV (CST, 11967 and 4850), GLUD (Novus, NBP1-68846), PDH (abcam, 110333), HADHA (abcam, 203114), Cox4-l1 (R&D, AF5814), Lamp1 (DSHB, H4A3-c), Calnexin (ThermoFisher, MA3-027), Golgin (ThermoFisher, 14-9767-82), Tom20 (Santa Cruz, sc-11415), Citrate Synthase (Sigma, SAB2702186). After washing with PBS, the samples were incubated with secondary antibodies for two hours at room temperature. All secondary antibodies used are conjugated with Alexa Fluor (Invitrogen) dyes—either 488, 555, or 647. Finally, the samples were washed and mounted onto glass slides using ProLong™ Diamond Antifade Mountant (Life Science) for imaging under widefield or confocal microscopes (Leica), as described in the figure.

### Correlation coefficient (r) measurement

Pearson’s correlation coefficient was measured using CellProfiler version 4.2.5 to assess the degree of overlap between GLS1 and mitochondrial protein staining. The obtained values for representative images are indicated in yellow in each figure.

### Mitochondrial fractionation and western blot

Mitochondrial subfractionation was performed by using the Mitochondria Isolation Kit (Thermo Fisher Scientific, 89874) following the manufacturer’s instructions. Mitochondrial fraction along with total cell lysate and cytosolic fractions were lysed in RIPA buffer that contained Complete mini protease inhibitor cocktail (Roche) and phosphatase inhibitor (PhosSTOP, Roche). Protein concentration was measured by BCA protein assay kit (Thermo Fisher Scientific) and the samples were then boiled in the Laemmli buffer and loaded into 4%–20% gradient gel (Bio-Rad), transferred to PVDF membrane (Millipore), and analyzed by immunoblotting. The following primary antibodies were used: GLS1 (abcam, 156876), Citrate Synthase (Sigma, SAB2702186), TIMM23 (abcam, 116329), HIF1a (Novus Bio, NB100-296), HIF2a (CST, 7096s), MYC-tag (CST, 2278s), mCherry (CST, 43590), 14-3-3 (CST, 8312), and GAPDH (CST, 5174). Secondary antibodies that are conjugated with HRP were purchased from CST. Signal was detected using the ECL system (ImageQuant LAS 4000, Amersham Biosciences, GE Healthcare) according to the manufacturer’s instructions.

### qPCR

mRNA isolation and cDNA synthesis were done by using the TurboCapture mRNA Kit (QIAGEN) according to the manufacturer’s instructions. qPCR was performed on the CFX384 Bio-Rad Real-Time PCR Detection System using SYBR Green. Sequences of the primers used in this study were as follows: *HIF1a* (forward, 5’-TATGAGCCAGAAGAACTTTTAGGC-3’, reverse, 5’-CACCTCTTTTGGCAAGCATCCTG-3’), *HIF2a* (forward, 5’-CTGTGTCTGAGAAGAGTAACTTCC-3’, reverse, 5’-TTGCCATAGGCTGAGGACTCCT-3’), *LDHA* (forward, 5’-TTGACCTACGTGGCTTGGAAG-3’, reverse, 5’-GGTAACGGAATCGGGCTGAAT-3’), *PGK1* (forward, 5’-GACCTAATGTCCAAAGCTGAGAA-3’, reverse, 5’-CAGCAGGTATGCCAGAAGCC-3’), *HK2* (forward, 5’-GGGACAATGGATGCCTAGATG-3’, reverse, 5’-GTTACGGACAATCTCACCCAG-3’), *ENO1* (forward, 5’-TGGTGTCTATCGAAGATCCCTT-3’, reverse, 5’-CCTTGGCGATCCTCTTTGG-3’), *ANGPT2* (forward, 5’-ATTCAGCGACGTGAGGATGGCA-3’, reverse, 5’-GCACATAGCGTTGCTGATTAGTC-3’), *PDK1* (forward, 5’-CTATGAAAATGCTAGGCGTCTGT-3’, reverse, 5’-TGGGATGGTACATAAACCACTTG-3’), *VEGFA* (forward, 5’-TTGCCTTGCTGCTCTACCTCCA-3’, reverse, 5’-GATGGCAGTAGCTGCGCTGATA-3’), *DLL4* (forward, 5’-TGGGTCAGAACTGGTTATTGGA-3’, reverse, 5’-GTCATTGCGCTTCTTGCACAG-3’), *VEGFR2* (forward, 5’-GGAACCTCACTATCCGCAGAGT -3’, reverse, 5’-CCAAGTTCGTCTTTTCCTGGGC-3’).

### Determination of media glutamine concentration

Metabolites were extracted from 100 µL of culture medium with 1 µL of 4 N hydrochloric acid, immediately followed by addition of 400 µL of dry-ice cold analytical-grade methanol. Each sample was spiked with 18.75 nmol of L-norvaline as an internal standard, provided in the methanol. Samples were incubated on dry ice for 15 min then centrifuged at 21,000 x g for 15 min at 4°C. Supernatants containing soluble metabolites were transferred to new tubes and dried under vacuum in a speedvac microcentrifuge concentrator. In a fume hood, dried samples were resuspended in 100 µL of room-temperature 1:1 (vol/vol) analytical-grade acetonitrile and MtBSTFA (N-methyl-N-(tert-butyldimetylsilyl)trifluoroacetamide) (Regis Technologies 1-270242-200), and were heated on a 70°C heat block for 90 min. Then samples were cooled to room temperature (about 5 min), centrifuged at 13,000 x g for 5 min, and the supernatant was transferred to GC-MS vials with polypropylene inserts for small volume samples. One microliter of the sample was injected via automatic liquid sampler (Agilent 7693A) into an Agilent 7890B gas chromatograph (GC) coupled with an Agilent 5977B mass selective detector (MSD) (Agilent Technologies). The GC was operated in splitless injection mode with helium as the carrier gas at a flow rate of 1.2mL/min. The GC column was a 30 m x 250 µm x 0.25 µm HP-5ms Ultra Inert column (Agilent 19091S-433UI). The inlet temperature was 250°C, and after 3 min at 100°C, the oven temperature program was increased as follows: 4°C/min to 230°C then 20°C/min to 300°C and hold 5 min. The transfer line temperature was 250°C, and the MSD source and quadrupole temperatures were 230°C and 150°C, respectively. After a 6 min solvent delay, the MSD was operated in electron ionization mode and scan mode with a mass range of 50-550 AMU at 2.9 scans/s. Agilent MassHunter Qualitative Analysis software (B.07.00) was used for visualization of chromatograms. A standard curve of glutamine in culture medium was used to determine glutamine concentrations.

### GLS1 activity assay

For the experiment in Figure 5, HUVECs were pre-conditioned in 2mM of Q vs noQ media for 24 hours. Subsequently, 2mM of uniformly labeled ^13^C glutamine (U-^13^C-Q) (Cambridge Isotope Laboratories) was introduced for the indicated time, and ^13^C glutamate was measured as a readout of GLS1 activity. Briefly, intracellular metabolites were extracted by aspirating the medium and quickly adding 80% methanol (MeOH) pre-chilled at -80°C. Following a 20-minute incubation on dry ice, the resultant mixture was scraped, collected into a tube, and centrifuged at 10,000 × g for 5 minutes. The supernatants were dried under nitrogen gas and analyzed using reversed-phase ion-pairing chromatography coupled with negative mode ESI high-resolution mass spectrometry on a stand-alone orbitrap. EI-MAVEN was used for peak picking and Accucore was used for natural isotope correction.

### LC/MS

Culture medium samples were centrifuged at 2000g for 5 min at 4°C to remove cell debris. 10 μl medium was extracted using 100 μl -20°C pre-chilled lysis buffer (40% methanol, 40% acetonitrile, 20% water). For analysis of cellular metabolites, cells were cultured in a 6-well plate and lysed using 200 μl pre-chilled lysis buffer. For LC/MS of tumors, frozen samples stored at -80°C were ground at liquid nitrogen temperature with a Cryomill (Retsch, Newtown, PA). Tissue powder was then weighed (∼20 mg) and extracted with pre-chilled lysis buffer (∼40x tissue weight, a concentration of 25 mg/ml). All LC/MS samples were then centrifuged twice at 16,000 g for 10 min at 4°C. The final supernatant was transferred to LC/MS tubes for analysis. Targeted measurements of glutamine, glutamate and a-ketoglutarate were achieved on a quadrupole orbitrap mass spectrometer (Thermo Fisher Scientific, Q Exactive) coupled to hydrophobic interaction chromatography (HILIC) via electrospray ionization. LC separation was performed on an XBridge BEH Amide Column (2.1 × 150 mm, 2.5 μM particle size and 130 Å pore size; Waters Corporation) using a gradient of solvent A (95:5 water: acetonitrile with 20 mM ammonium acetate and 20 mM ammonium hydroxide, pH 9.45) and solvent B (acetonitrile). The gradient was: 0 min, 90% B; 2 min, 90% B; 3 min, 80% B; 5 min, 80% B; 6 min, 75% B; 7 min, 75% B; 8 min, 70% B; 9 min, 70% B; 10 min, 50% B; 11 min, 50% B; 12 min, 40% B; 14 min, 40% B; 15 min, 90% B; 20 min, 90% B. The injection volume was 5 μl and the autosampler temperature was set at 4°C. The total running time was 20 minutes at a flow rate of 150 μl/min. The data was generated using negative ion mode with a scan range of 65-835 m/z and resolution of 140,000, and normalized by cell number and total ion counts.

### Mitochondrial fragmentation count (MFC) as a mitochondrial fission index

MFC was calculated by counting non-contiguous mitochondrial particles and dividing by the number of pixels which comprise the mitochondrial network^48^.

### WT vs K320A GLS1 overexpression

WT or K320A GLS1 was overexpressed in HeLa, UMRC2, and 769-P cells using a retroviral infection system. GLS1-myc-mCherry sequences were cloned into the retroviral pLHCX plasmid (Addgene, 44239) and transfected to HEK293T cells for virus generation. Virus-containing media was filtered and used for transduction of HeLa, UMRC2, and 769-P with WT or K320A GLS1.

### Human kidney sample immunohistochemistry

Kidney cancer tissue microarray slides (T071b) are purchased from TissueArray. The slides were deparaffinized in xylene, rehydrated in a series of ethanol solutions, and quenched in 0.3% hydrogen peroxide/methanol for 15 min. For antigen retrieval, slides were boiled for 20 min in 10 mM sodium citrate (pH 6.0). Sections were blocked with 5% goat serum/1% BSA/0.5% Tween-20 for 1 h and incubated with primary antibodies diluted in blocking buffer overnight at 4 °C. Following primary antibody, slides were incubated with biotinylated secondary antibodies for 1 h followed by ABC solution (Vector Laboratories) for 30 min. The slides are then developed with 3,3 ′-diaminobenzidine (Vector Laboratories). Slides were counterstained with hematoxylin, dehydrated, and mounted with Permount (ThermoFisher Scientific). Primary antibodies used: GLS (1:200; Abcam; Cat. #ab156876).

### Tumorigenesis assay

Female NIH-III nude mice (Charles River Laboratories, strain code: 201) that were 6-8 weeks old were subcutaneously injected in each flank with ten million UMRC2 cells overexpressing either WT-GLS1 or K320A-GLS. Each mouse received both a WT and K320A GLS1 tumor on separate flanks, ensuring each mouse served as its own control for a pair matched comparison. Cells were resuspended in ice-cold PBS and combined 1:1 with Matrigel (BD Biosciences, 356234) for a final volume of 200*μ*L per injection. Tumor volumes were recorded at the indicated time points using caliper measurements, calculated by the formula V=(π/6)(L)(W2), where L was the longer measurement and W was the shorter measurement. Tumors were harvested at the 12-week time point for weight and immuno-histochemistry analysis. Following embedding in OCT (Sakura), tumor section and staining were performed as previously reported^49^. In brief, tumors were sectioned at 5 μm. Slides were treated with 0.2% Triton X-100 in PBS and incubated in blocking solution (1% BSA, 5% goat serum in PBS) for 1 hour at room temperature. Primary antibodies anti-mcherry (abcam, ab205402, 1:200) and anti-COX4 (CST, 11967S, 1:100) were used for overnight incubation at 4℃, followed with staining of secondary antibodies (Invitrogen, Alexa-Fluor 555 conjugated and Alexa-Fluor 488 conjugated, 1:400) or TUNEL enzymatic mixture (Roche, 11684795910) according to manufacturer’s instructions. Slides were mounted using ProLong gold antifade reagent with DAPI (Invitrogen, P36935) and examined by Zeiss LSM710 confocal microscope. TUNEL/mcherry double positive cells were quantified manually in each image for apoptosis analysis.

### Statistics

Statistical comparisons among study groups were performed using either a 2-tailed Student’s t-test or 1-way ANOVA, followed by Bonferroni’s post hoc testing. A significance level of P < 0.05 was considered statistically significant. All data are presented as mean ± SD. Results from cell culture experiments are representative of a minimum of 3 independent experiments.

## Study approval

All mouse experiments were performed according to procedures approved by the University of Pennsylvania Institute for Animal Care and Use Committees (Philadelphia, PA).

## Author contributions

BK and WZ led the studies and were directly involved in most experiments. BK and ZA oversaw the studies. WZ, BK, NJC, SB, YJ, CB, MN, CJ, and BK conducted experiments and acquired data. WZ, CS, ZA, and BK interpreted data. WZ, ZA, and BK wrote the manuscript. All authors discussed the results and commented on the manuscript.

## Acknowledgements

Boyoung Kim was supported by Basic Science Research Program through the National Research Foundation of Korea (NRF), funded by the Ministry of Education (RS-2024-00412498). NJC was supported by NCI (F30CA271654). MN was supported by NCI (F31CA261041). CJ was supported by NIAAA (AA029124). ZA was supported by NHLBI (HL167014), the DOD (KC220099), and the Ludwig Foundation. Boa Kim was supported by NHLBI (R56HL162660) and AHA (24CDA1264317). We thank the Microscopy Services Laboratory (UNC) for assistance with confocal imaging. The Microscopy Services Laboratory, Department of Pathology and Laboratory Medicine, is supported in part by P30 CA016086 Cancer Center Core Support Grant to the UNC Lineberger Comprehensive Cancer Center.

---

**SFigure 1.**
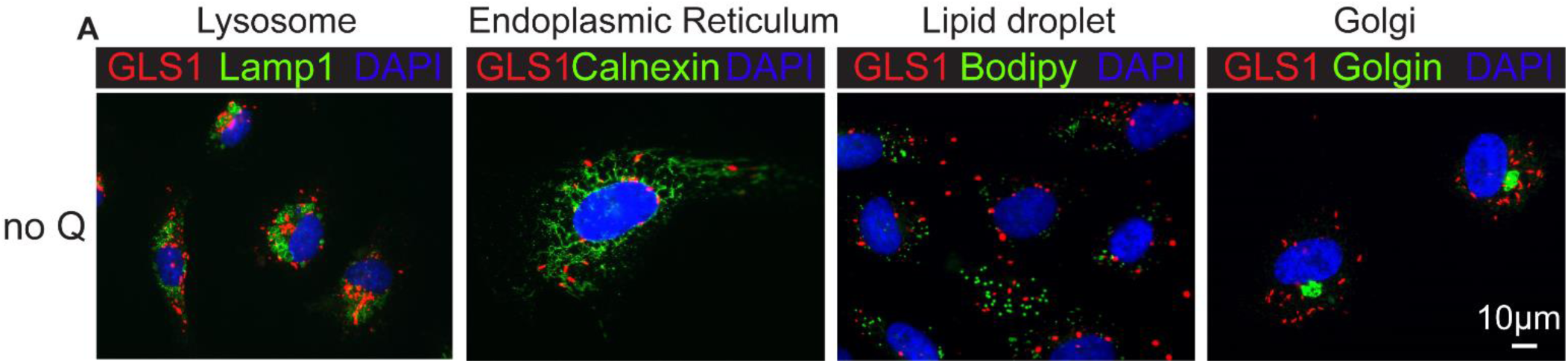
Clustered GLS1 does not co-localize with non-mitochondrial organelles. Co-staining of GLS1 with markers of various intracellular organelles in HUVECs upon 24-hour culture in noQ media: lysosome (Lamp1), endoplasmic reticulum (Calnexin), lipid droplet (BODIPY), or golgi (golgin).

**SFigure 2.**
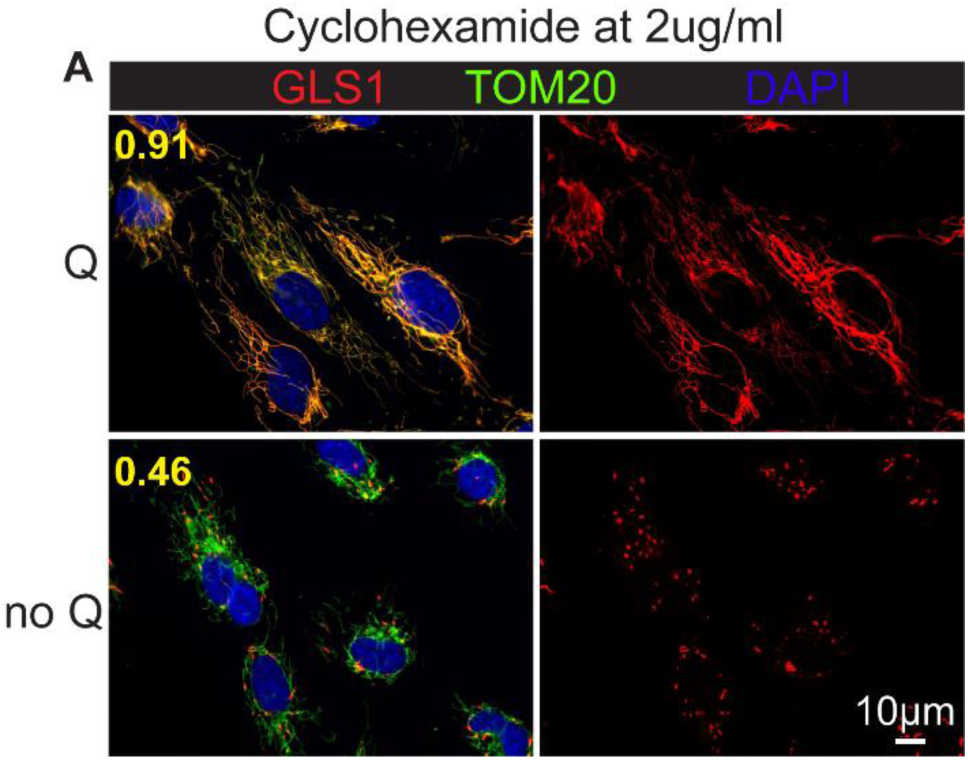
Existing, but not the newly synthesized, GLS1 cluster upon glutamine deprivation. ICC of GLS1 (red) and TOM20 (green) in HUVECs after a 24-hour culture in Q vs. noQ media in the presence of cycloheximide (2μg/ml).

**SFigure 3.**
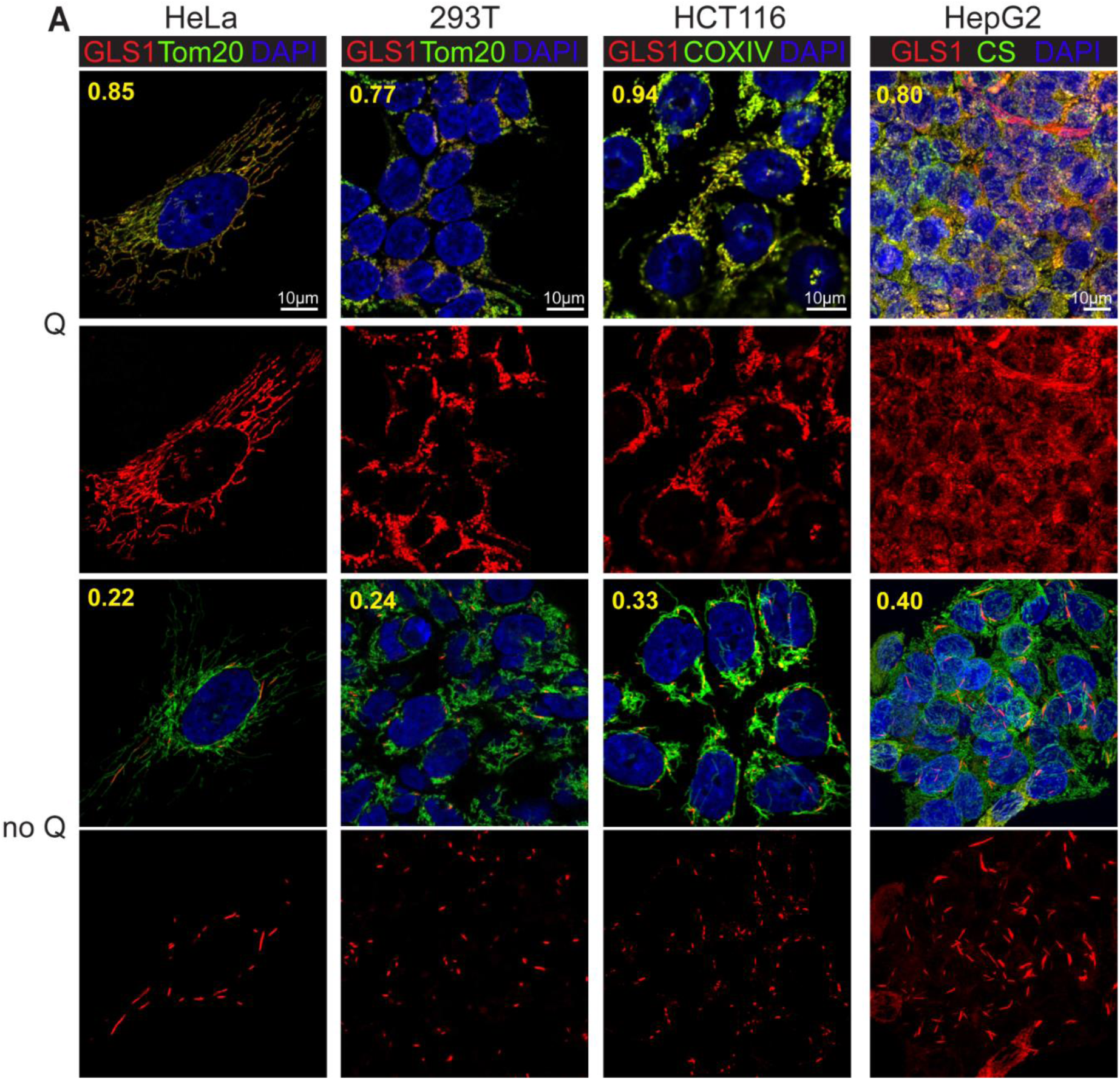
Glutamine deprivation-induced GLS1 clustering occurs ubiquitously in various cell types. CC of GLS1 (red) and mitochondrial marker proteins (Tom20, COXIV, or CS in green) in various cell types: HeLa, 293T, HCT116 and HepG2.

**SFigure 4.**
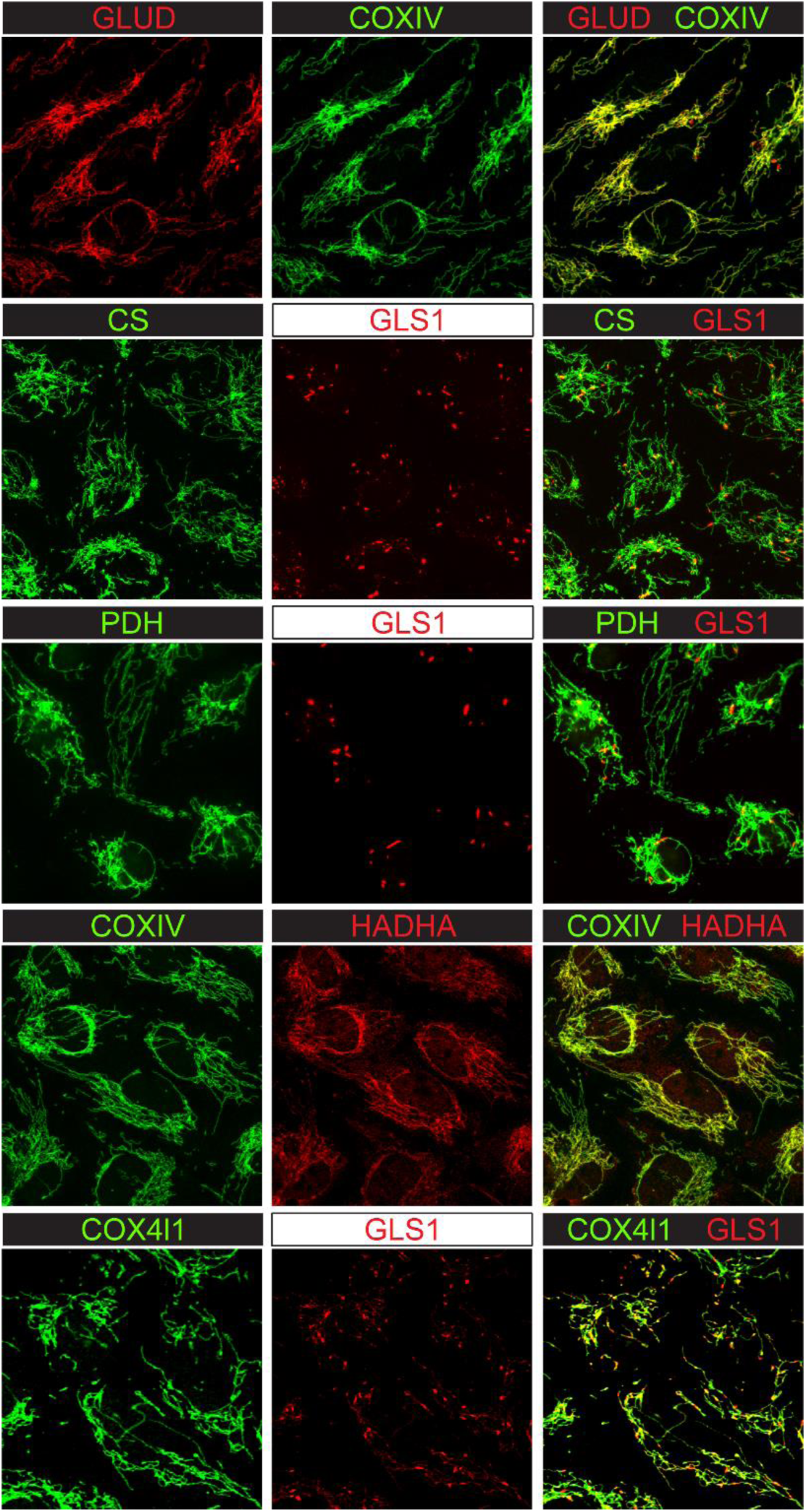
Clustering upon glutamine deprivation occurs uniquely to GLS1. ICC of various mitochondrial proteins in HUVECs after 24-hour culture in noQ media: GLUD, COXIV, CS, PDH, HADHA, and COX4l1.

**SFigure 5.**
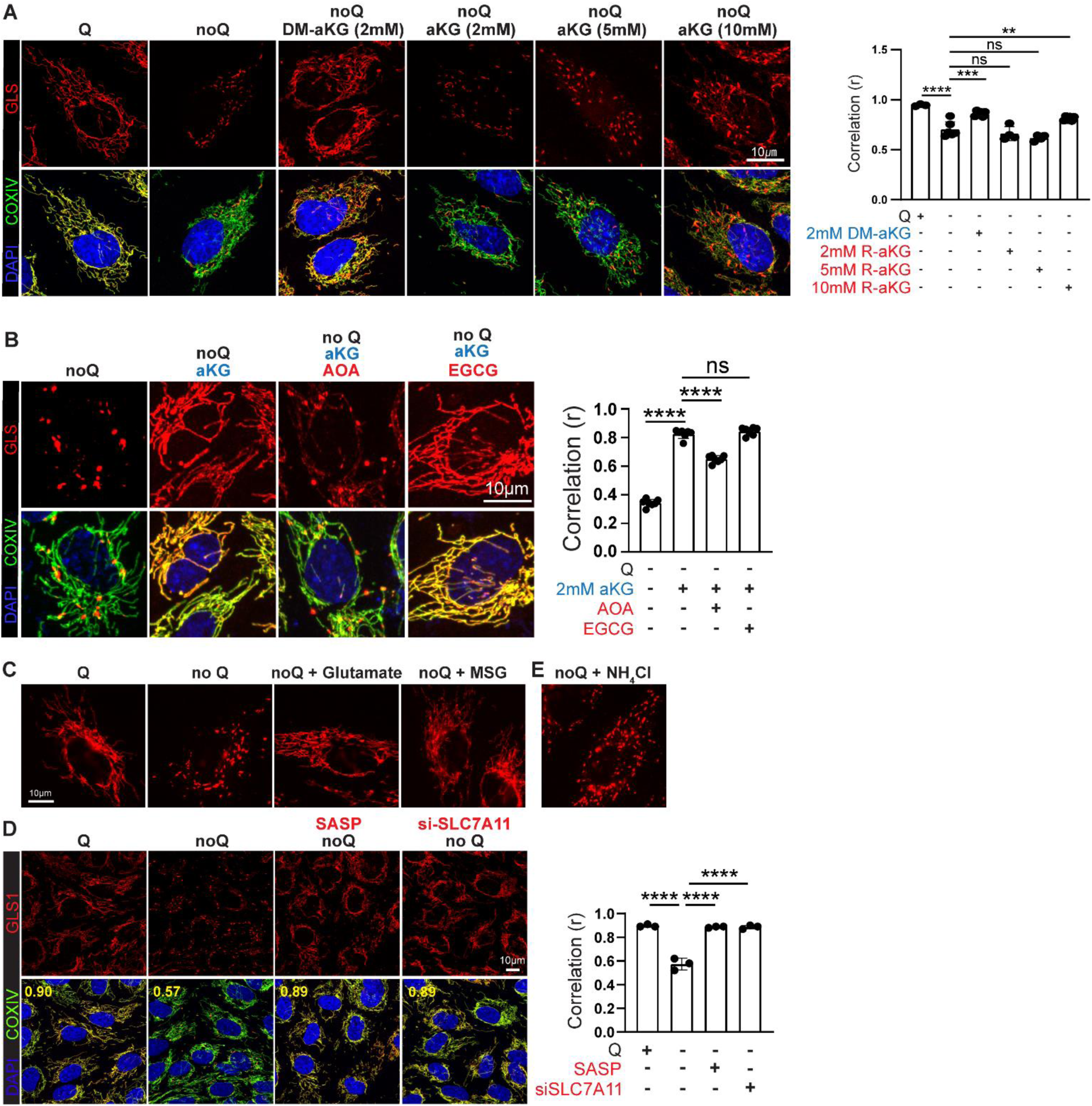
Restoration of glutamate level restores GLS1 clustering. **A**. ICC demonstrating the rescued GLS1 clustering by DM-aKG and aKG in HUVECs. ** p < 0.01, *** p < 0.001, **** p < 0.0001 and ns (not significant, p ≥ 0.05) by 1-way ANOVA. **B**. ICC showing the reversal of the aKG-induced GLS1 redistribution by AOA or EGCG. **** p < 0.0001 and ns by 1-way ANOVA. **C**. Rescued GLS1 clustering by the supplementation of glutamate or monosodium glutamate (MSG) in noQ for 6 hours. **D**. Rescue of GLS1 clustering by chemical inhibition (by SASP treatment at 300μM) or siRNA knockdown of SLC7A11. **** p < 0.0001 by 1-way ANOVA. **E**. No effect of supplementation of ammonia by NH_4_Cl on GLS clustering.

**SFigure 6.**
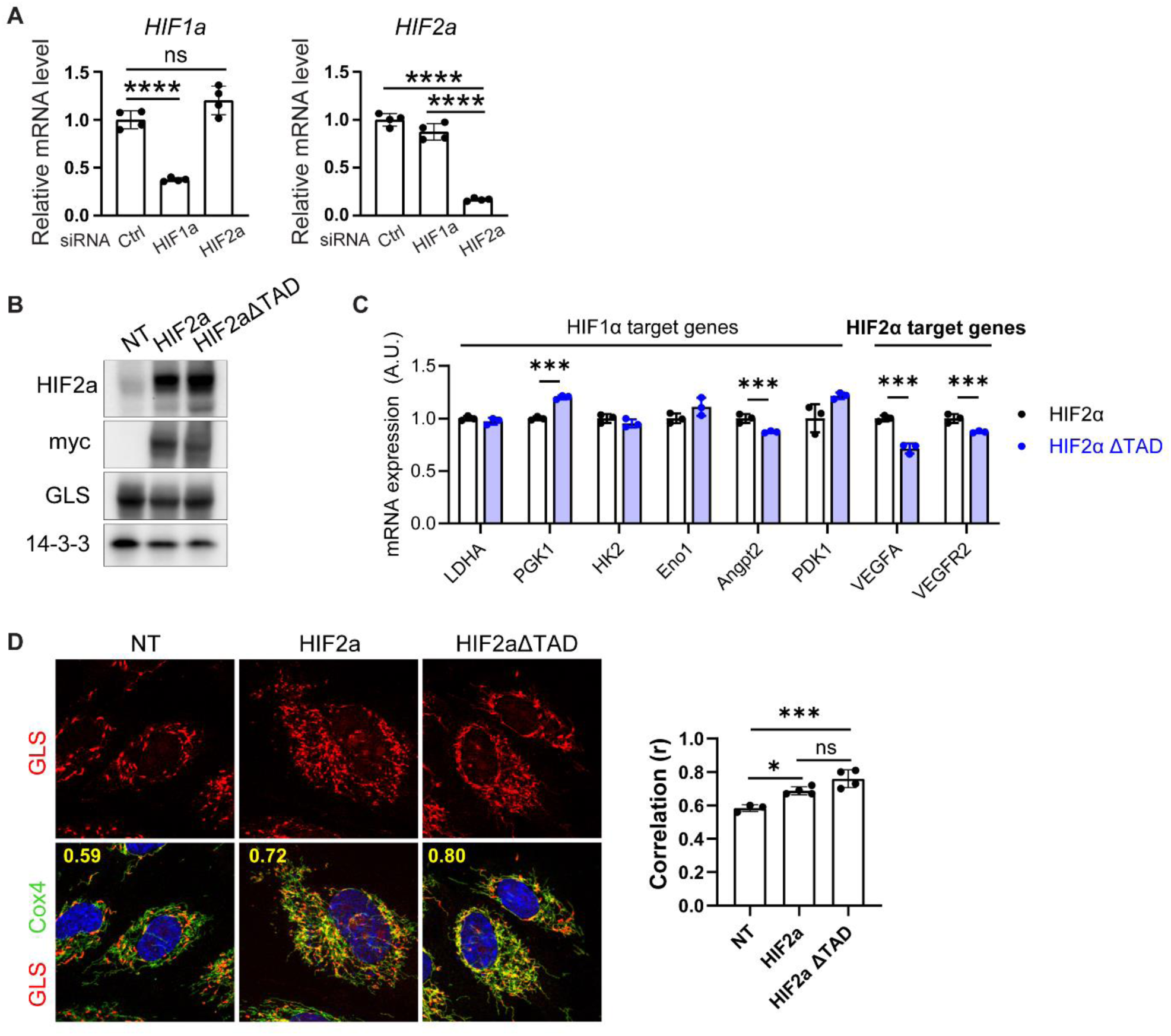
HIF2a protein but not its transcriptional activity inhibits GLS1 clustering. **A**. qPCR analysis showing the validation of si_HIF1a and si_HIF2a in HUVECs. **** p < 0.0001 and ns by 1-way ANOVA. **B**. Validation of the overexpression of HIF2a and HIF2aΔTAD in HUVECs by western blotting. **C**. qPCR analysis showing the suppression of HIF2aΔTAD on the mRNA expression of HIF2a target genes. *** p < 0.001 by 1-way ANOVA. **D**. ICC showing the inhibition of GLS1 clustering by the overexpression of HIF2a and HIF2aΔTAD. * p <0.05, *** p < 0.001 and ns by 1-way ANOVA.

**SFigure 7.**
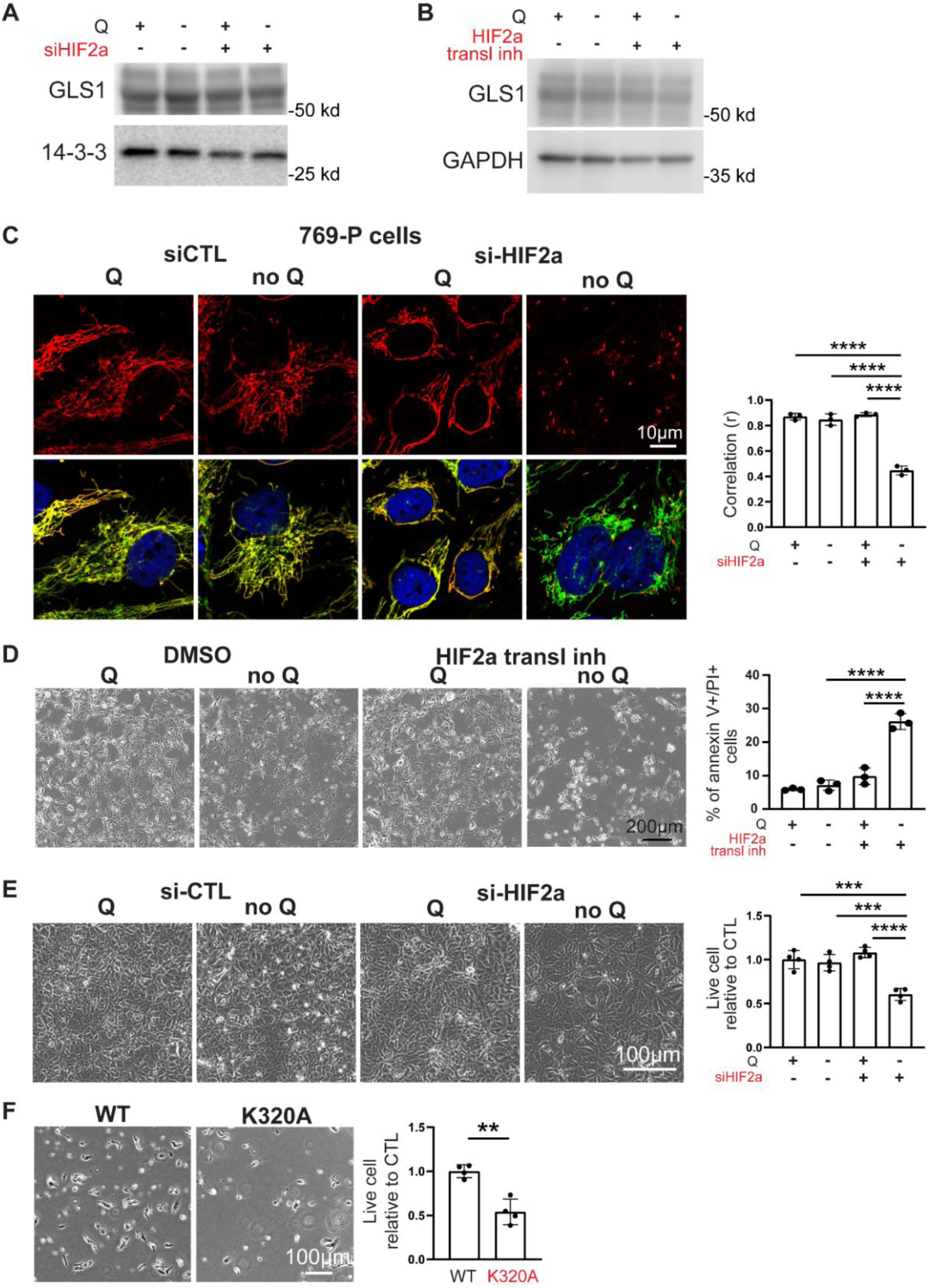
GLS1 clustering is prevented in 769-P cells in a HIF2α-dependent manner. **A** and B. Western blotting analysis showing the effects of HIF2a siRNA (A) and its translational inhibitor (B) on the expression of GLS1 in UMRC2 cells. **C**. Resistance to noQ-induced GLS1 clustering in 769-P cells is reversed by si-HIF2α. **** p < 0.0001 by 1-way ANOVA. **D**. Resistance to noQ-induced cell death in 769-P cells is reversed by treatment with an inhibitor of HIF2α translation. **** p < 0.0001, *** p < 0.001 by 1-way ANOVA. **E**. Resistance to noQ-induced cell death in 769-P cells is reversed by siHIF2α. **** p < 0.0001, *** p < 0.001 by 1-way ANOVA. **F**. Increased cell death in 769-P cells overexpressing the K320A mutant GLS1 compared to those overexpressing WT GLS. *** p < 0.001 by t-test.

**SFigure 8.**
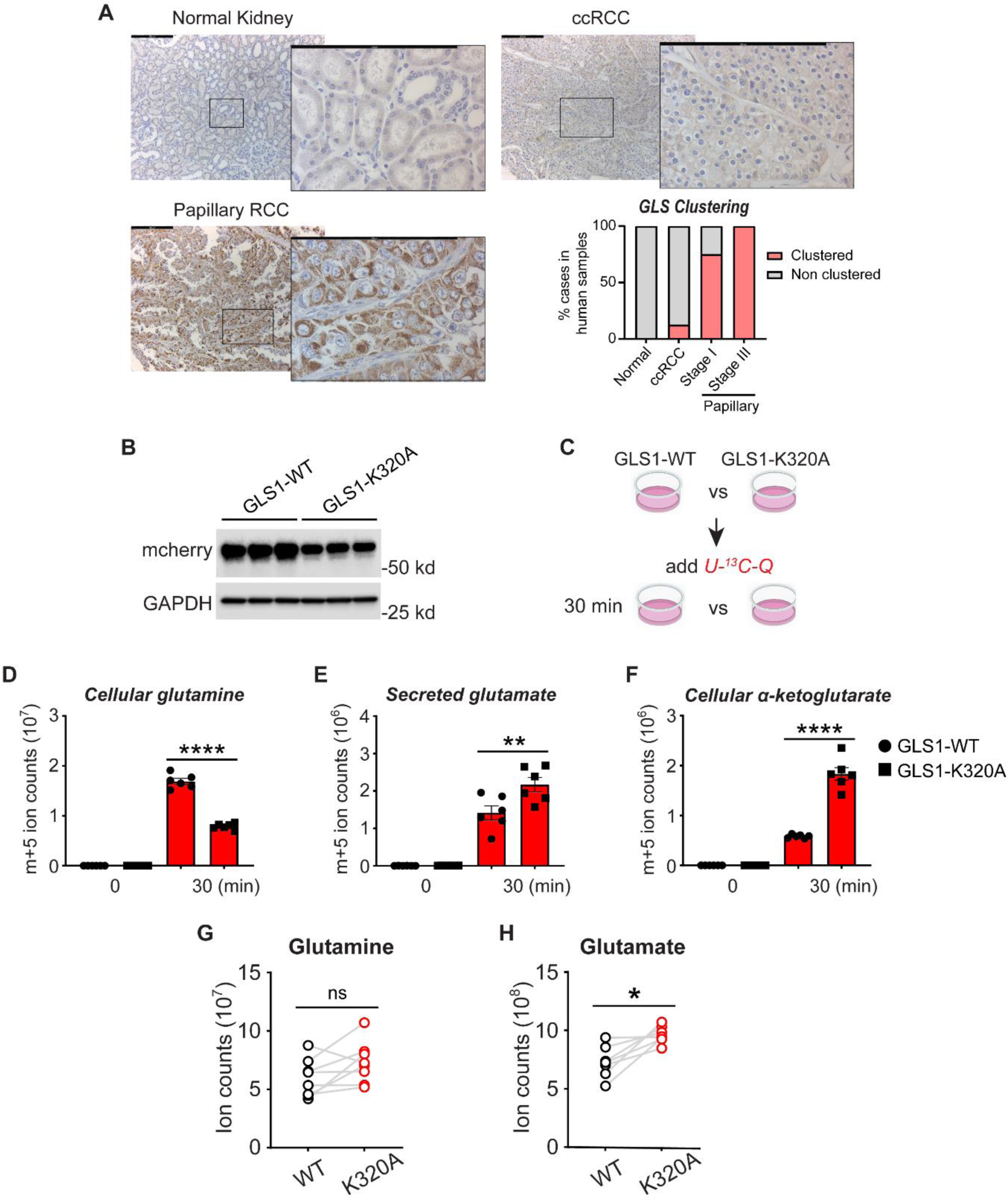
Enhanced enzymatic activity of GLS-K320A vs. GLS-WT in UMRC2 cells. **A**. Reduced GLS1 clustering in human ccRCC samples compared to Papillary RCC. Images show the IHC of GLS1 in human clinical samples. Magnifications of the area in the block are shown on the right of each image. For normal kidney and ccRCC, 8 areas from 2 independent samples were quantified. For Papillary RCC stage I and stage III, 4 areas from one independent sample were quantified. Scale bars are 200 μm. **B**. Western blotting analysis showing the validation of the expression of GLS1-WT and GLS1-K320A in UMRC2 cells. **C**. Experimental scheme of U-^13^C-Q tracing assay in UMRC2 cells. **D**-**F**. Quantifications of m+5 metabolites including glutamine (D), glutamate (E) and aKG (F) with conditions shown in C. ** p < 0.01 and **** p < 0.0001 by 1-way ANOVA. **G** and **H**. Quantifications of glutamine (G) and glutamate (H) in UMRC2 tumors. * p < 0.05 and ns by paired t-test.

